# Evaluating possible maternal effect lethality and genetic background effects in *Naa10* knockout mice

**DOI:** 10.1101/2023.04.27.538618

**Authors:** Gholson J. Lyon, Joseph Longo, Andrew Garcia, Fatima Inusa, Elaine Marchi, Daniel Shi, Max Dörfel, Thomas Arnesen, Rafael Aldabe, Scott Lyons, Melissa A. Nashat, David Bolton

## Abstract

Amino-terminal (Nt-) acetylation (NTA) is a common protein modification, affecting approximately 80% of all human proteins. The human essential X-linked gene, *NAA10,* encodes for the enzyme NAA10, which is the catalytic subunit in the N-terminal acetyltransferase A (NatA) complex. There is extensive genetic variation in humans with missense, splice-site, and C-terminal frameshift variants in *NAA10*. In mice, *Naa10* is not an essential gene, as there exists a paralogous gene, *Naa12*, that substantially rescues *Naa10* knockout mice from embryonic lethality, whereas double knockouts (*Naa10*^−/Y^ Naa12*^−/−)^* are embryonic lethal. However, the phenotypic variability in the mice is nonetheless quite extensive, including piebaldism, skeletal defects, small size, hydrocephaly, hydronephrosis, and neonatal lethality. Here we replicate these phenotypes with new genetic alleles in mice, but we demonstrate their modulation by genetic background and environmental effects. We cannot replicate a prior report of “maternal effect lethality” for heterozygous *Naa10^−/X^* female mice, but we do observe a small amount of embryonic lethality in the *Naa10^−/y^*male mice on the inbred genetic background in this different animal facility.

## INTRODUCTION

Targeting 40% of the human proteome, NatA, the major N-terminal acetyltransferase (NAT) complex, acetylates <u>Ser</u>-, <u>Ala</u>-, <u>Gly</u>-, Thr-, Val-, and Cys-N-termini following cleavage of the initiator methionine [1,2]. The canonical human NatA consists of two main subunits, the catalytic subunit N-α-acetyltransferase 10 (*NAA10*) (Ard1) and the auxiliary subunit NAA15 (Nat1) and engages with a regulatory subunit, HYPK [3–5]. N-terminal (Nt-) acetylation (NTA) is one of the most common protein modifications, occurring co- and post-translationally [6,7]. Approximately 80% of cytosolic proteins are N-terminally acetylated in humans and ∼50% in yeast [1], while NTA is less common in prokaryotes and archaea [6].

NTA is catalyzed by a set of enzymes, the NATs, which transfer an acetyl group from acetyl coenzyme A (Ac-CoA) to the free α-amino group of a protein’s N-terminus. To date, eight distinct NATs (NatA – NatH) have been identified in metazoan (NatA-F and NatH) and plant (NatG) species that are classified based on different conserved subunit compositions and substrate specificities [2,7,8]. NTA has been implicated in steering protein folding, stability or degradation, subcellular targeting, and complex formation [9–13]. Particularly, Naa10-catalyzed N-terminal acetylation has been reported to be essential for development in many species [14–18] and although NatA is not essential in *S. cerevisiae*, depletion of *Naa10* or Naa15 has strong effects, including slow growth and decreased survival when exposed to various stresses [19,20]. In addition, it has been recently shown that mice have a compensating enzyme Naa12, which prevents embryonic lethality in the *Naa10* knockouts [21], but a similar gene has not been found in humans. Furthermore, given that *NAA10* was also identified in screens for essential genes in human cell lines [22,23], it seems unlikely that an unknown *NAA10*-like paralogous gene exists in humans, other than the already known *NAA11* [24,25].

Ogden syndrome (OS), was first reported and named in 2011 after the location of the first affected family residing in Ogden, Utah, USA [26,27]. In that first family, five males were born to four different mothers and all the males died in early infancy, with a range of cardiac, phenotypic variation and other defects, including lethal cardiac arrhythmias. The underlying genetic defect was characterized as a single missense change coding for Ser37Pro in the X-linked gene, *NAA10*, and was confirmed in a second independent family in California, USA, with three males that also died during infancy. The identical variant was recently reported in a third family [28], and a fourth family [29]. There is a *S. cerevisiae* model for the *Naa10* Ser37Pro mutant, in which that variant impairs NatA complex formation and leads to a reduction in both NatA catalytic activity and functionality [30,31]. Furthermore, OS patient-derived cells have impaired *in vivo* NTA of a few NatA substrates [32].

Since the initial discovery of OS in 2011, multiple groups have reported additional variants either in *NAA10* in both males and females [33–46] or in the heterodimeric protein partner encoded by *NAA15* [35,47–50]. The genetic landscape of variation in *NAA10* and *NAA15* in humans has been presented recently with many more cases of Ogden syndrome (OS) (also known as “*NAA10*-related neurodevelopmental syndrome”), and “*NAA15*-related neurodevelopmental syndrome” [51].

Previous publications have reported that one of the phenotypes observed in *Naa10* knockout mice is low body weight compared to their wildtype littermates [21,52]. A recent analysis in humans analyzed these growth defects in greater detail, which showed extensive weight fluctuations, along with the recommendation that OS individuals not tracking above the failure to thrive range past one year of age might be considered for G-tube placement to avoid prolonged growth failure [53]. The current study aims to characterize the development, growth and phenotypes of various mice deficient in *Naa10*, beginning with embryogenesis, thus extending and expanding on our prior findings [21].

## RESULTS

### Independently generated knockdown (KD) and knockout mice for *Naa10* demonstrate similar phenotypes

Twice we attempted to develop a mouse model for Ogden Syndrome with a Ser37Pro missense mutation found in the first family identified with this syndrome [26]. The first attempt included the use of a minigene for exons 2-3, which was inserted in an intron in *Naa10*, that could theoretically then be activated upon inversion of the cassette by crossing to Cre-expressing animals (see **S1 Fig** for gene targeting design). Unfortunately, both Western blotting and quantitative PCR revealed that the insertion of this minigene in the intron between exons 3 and 4, regardless of its inversion status, led to >95% knockdown of expression of *Naa10* (**S2 and S3 Figs**). As such, the interpretation of any results in this mouse model for the Ogden Syndrome Ser37Pro mutation are complicated by the knockdown of *Naa10*. We refer to these mice as male *Naa10^mini/Y^*, female *Naa10^mini/mini^*, male *Naa10^invS37P/Y^*, and female *Naa10 ^invS37P/invS37P^*. When tabulating the Mendelian ratios, with genotyping performed after weaning around four weeks of age, there was a slight deviation (22%) from the predicted Mendelian ratio (25%) for the male *Naa10^mini/Y^* mice (**S1 Table**) and a much larger deviation (13%) for the male *Naa10^invS37P/Y^* mice (**S2 Table**), which suggests the possibility that the S37P mutation does indeed exert a hypomorphic effect on NatA function. However, the confound for any interpretation of these data is the presence of severe knockdown of overall *Naa10* protein expression, given that the minigene apparently has some effect on RNA expression or stability (**S2B Fig**).

The phenotypes of these Naa10 knockout mice are similar to previously published identifying characteristics including piebaldism, hydrocephaly, cardiac defects, homeotic anterior transformation, and urogenital anomalies [21]. Although there was complete penetrance for piebaldism in the male mice, there was variable amounts of this, which was quantified to show the high variability of this phenotype in this allelic series of mutant mice (**S4 Fig**). Piebaldism was also present and quantified in female *Naa10^mini/mini^* and female *Naa10 ^invS37P/invS37P^* mice, with no obvious correlation between the amount of piebaldism and the genotype, age, or weight of the mice (**S5 Fig**). Piebaldism in heterozygous females was only very rarely seen (on the order of 1-2 animals among >40 animals per allele). Another phenotype identified with complete penetrance was bilateral supernumerary ribs (14 pairs of rib instead of 13) in all male *Naa10^mini/Y^*, female *Naa10^mini/mini^*, male *Naa10^invS37P/Y^*, and female *Naa10 ^invS37P/invS37P^* (**S3 Table**). This extra pair of ribs linking to the sternum transforms the T8 vertebrae into an anterior T7-like phenotype. Many of these mice also had four instead of the usual three sternebrae, which were sometimes fused. Cervical vertebrae fusion was also demonstrated in these mice, particularly involving C1 and C2, suggesting possible anteriorization of C2 into a C1-like phenotype (**S4 Table**). These phenotypes are identical to what was described in the *Naa10^−/y^* mice [21].

Bone density of calvarias was measured using computerized tomography (CT) scanning (**S6 Fig)**, showing no difference from wild type, except in a few of the *Naa10^−/y^* mice where hydrocephaly developed, accompanied by dilatation of the skull over time with thinning of the calvarium. As such, the published calvarial bone density phenotype reported in three-day old *Naa10^−/Y^* mice [54] does not remain by adulthood. Femur bone density did show a small but statistically significant decrease in the male *Naa10^−/y^* mice and *Naa10 ^invS37P/y^* mice, compared to the *Naa10^+/y^* and *Naa10^mini/y^* mice (**S7B Fig**), and the female heterozygous *Naa10^+/invS37P^* also had slightly decreased femur bone density compared to *Naa10^+/+^* and *Naa10^+/−^*mice (**S7D Fig**). However, these data are limited by the fact that these measurements were taken in mice at all ages, and the overall number of mice was small in some groups (e.g., n=3 for *Naa10^+/invS37P^*) (**S7A** and **S7C Fig**). Furthermore, the *Naa10^−/Y^* mice were still on a somewhat mixed genetic background at the time of these experiments, as the number of backcrosses to C57BL/6J had only reached about 12-13 backcrosses at that time. Future experiments should ideally repeat this with a larger number of mice at one age timepoint, ideally on an inbred C57BL/6J genetic background (>20 backcrosses or mice generated at the outset from zygotes from an inbred background).

The second attempt to generate *Naa10 ^S37P/y^* mice included four rounds of microinjection (see **S5 Table**) of CRISPR reagent mix including guide RNA and oligonucleotide donor into zygotes obtained from the mating of B6D2F1 females (i.e., 50% C57BL/6J, 50% DBA/2J (D2)) females to inbred C57BL/6J males (Jackson Laboratories, Bar Harbour, ME, USA). The guide RNA was produced and validated from Horizon (Perkin Elmer, USA), using a Cel1-nuclease assay, and the most active guide was selected, including the targeting cr-RNA sequence and the tracrRNA portion. Despite screening 156 offspring from these four injections, only indels were obtained, with no evidence of homologous recombination with the oligonucleotide donor to generate the desired Ser37Pro missense mutation. Two of the indels, namely a one base pair deletion (Δ668) and a seven base pair deletion (Δ668-674) were successfully transmitted to the next generation by backcrossing to C57BL/6J mice. Western blotting and qPCR confirmed knockout of *Naa10* due to the frameshifts introduced by the indels (**S8 Fig**). Breeding of these mice on this mixed genetic background did not show any obvious embryonic or neonatal lethality for the Δ668 male mice or the Δ668-674 male mice, as they were born and genotyped in the first week of life with no deviation from the expected 25% Mendelian ratios (**S6** and **S7 Tables**). Although most of these knockout mice had piebaldism, some of them did not (6 out of 81 Δ668 male mice and 10 out of 36 Δ668-674 male mice without piebaldism). The *Naa10* indel mice also have bilateral supernumerary ribs (14 pairs of rib instead of 13), and a majority of these mice also had four instead of the usual three sternebrae, which were sometimes fused (**S3 Table**), as previously reported [21]. Cervical vertebrae fusion was also demonstrated in these mice (**S4 Table**). Out of all mice that were generated in the above and other matings, hydrocephaly did develop in the *Naa10* Δ668 male mice (23/62, or 37%) and in the *Naa10* Δ668-674 male mice (16/45, or 36%), which is similar to the previously reported rate around 40% [21] and substantially higher than the rate of 1% in wild type mice on the C57BL/6J genetic background.

As these new strains of knockdown or knockout mice recapitulated the phenotype of the already established *Naa10* knockout (KO) lines [21], it was decided to not maintain these lines, so the minigene and indel strains were euthanized, with sperm cryopreservation at Cold Spring Harbor Laboratory (CSHL), available upon request.

### Validation of a specific *Naa10* antibody and demonstration of heterozygous expression in female *Naa10* mice

Prior Western blotting using the *Naa10* antibody obtained from Abcam always demonstrated a cross-reactive band of the same molecular weight as *Naa10*, which can be seen in the Western blot data in **S2** and **S3 Figs**. We hypothesized that this cross-reactive band might be the recently discovered mouse Naa12 [21]. A new rabbit monoclonal anti-*Naa10* antibody is now available from Cell Signaling Technologies, Danvers, MA, USA, #13357, and this antibody was made with synthetic peptide corresponding to residues surrounding Asp204 of human NAA10. This is a region that diverges from mouse Naa12 [21]so, there should not be any cross-reactivity to Naa12. This was confirmed by Western blotting from tissues isolated from male mice. Biological replicates (n = 8) were obtained consisting of *Naa10^−/Y^* (n = 4) and *Naa10^+/Y^*(n = 4) animals **(S9 Fig**). Per genotype, the mean *Naa10* signal normalized to total protein detected in *Naa10*^−/Y^ was <10% of the *Naa10*^+/Y^ animals, a significant difference between genotypes (2-way ANOVA, F statistic = 14.52 on 3 and 12 DF, *P < 0.05).

Using this *Naa10*-specific antibody, we quantified the relative amounts of *Naa10* in heterozygous female mice (**S10** and **S11 Figs**). For quantification, Naa10 signal was normalized to total protein in each lane of a gel; post-transfer membranes were stained for total protein and scanned to verify transfer and equal loading before proceeding with Naa10 immunostaining. Liver and heart tissue lysates were prepared from *Naa10^−/+^*and/or *Naa10^+/−^* (n = 3) and *Naa10*^+/+^ (n = 3) mice obtained from our animal colony. The former heterozygous genotypes differ based on the *Naa10* knockout allele’s parent-of-origin. Based on the lack of maternal or paternal effect lethality in the embryonic and postnatal data (discussed below), we grouped the two *Naa10* heterozygous mutant genotypes together. Western blots using samples from these were repeated in triplicate. Replicate analysis indicates the mean normalized Naa10 signal in *Naa10*^+/−^ or *Naa10*^−/+^ animals is approximately 38% that of *Naa10^+/+^* animals, a statistically significant difference (2-way ANOVA, F-statistic = 12.52 on 3 and 32 DF, *P < 0.05, R-Studio, Boston, MS, USA). According to *post hoc* Tukey testing, the main source of variation in mean NAA10 signal is attributable to genotype differences; organ type is not a significant source of variation, though it participates in a significant interaction with genotype. Notably, the reduction in normalized *Naa10* signal was more variable compared to the *Naa10*^−/Y^ dataset. However, for the *Naa10*^−/Y^ dataset, one biological replicate was examined once in each organ, whereas each organ sample in the heterozygous mutant dataset was replicated three times. An earlier experiment **(S11 Fig)** used liver lysates from a separate set of mice (n = 6) composed of *Naa10^−/+^* (n = 3) and C57 females (n = 3); the mutant line [54] is now genetically inbred, with over 20 backcrosses to C57BL/6J. Compared to wild-type liver lysate, *Naa10* levels are substantially reduced (>50%) in heterozygous liver, though this is a non-significant result due to variability (Welch’s 2-sample t-test, t = −1.6784, df = 9.1762, P = 0.1252). Heterozygous female mice can undergo random X-chromosome inactivation, which can lead to Naa10 variability within and between tissues.

### Effects of environment, genetic background, and parent-of-origin

Given our findings related to genetic background on the phenotypes in the mice discussed above, we analyzed the phenotypes of *Naa10* mice after being backcrossed for 20 generations with C57BL/6J mice; however, this analysis was complicated by the change of colony venue in March 2019, when the backcross was at the 15th generation. Once the backcross was completed to 20 generations, the embryonic dissections and live births occurred in this new environment. A second aspect of the experiment was meant to address whether *Naa10* is associated with “maternal effect lethality”, as a different group argued that maternal inheritance of the *Naa10* knockout allele can have an effect with possible embryonic lethality for ∼20% of heterozygous Naa10^−/X^ females [52]. They used a nomenclature where Naa10^−/X^ mice inherit the null allele maternally, whereas *Naa10*^X/−^ female mice inherit the null allele paternally. We will use Naa10^−/X^ and Naa10^−/+^ interchangeably in the remainder of this manuscript. We have previously presented data that minimizes this effect and suggested lethality might be due to other unidentified factors, such as decreased maternal care of offspring [21]. To address this further, several new breedings were undertaken, as detailed in **Fig 2A** and **2B**. For example, embryos with maternal grandfather inherited *Naa10* knockout are designated as the AAA mating series, whereas embryos with maternal grandmother inherited *Naa10* knockout alleles are designated as the DDD mating series (with these letter combinations assigned based on internal lab databases).

**Fig 1.**
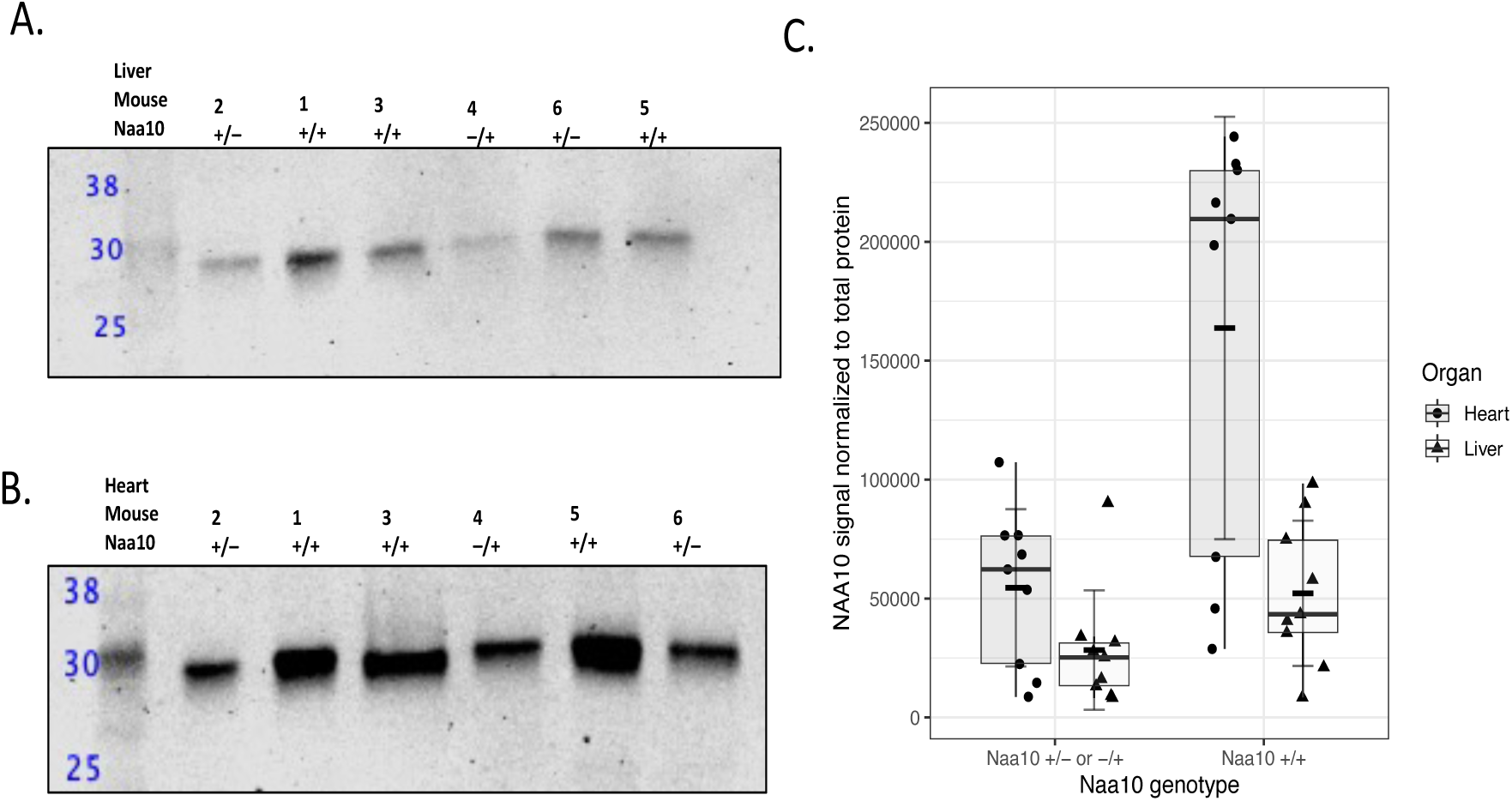
Western blot analysis of liver and heart lysates from *Naa10* heterozygous mutant female mice. Biological replicates (n = 6) were obtained from *Naa10*^+/–^ or *Naa10^−/+^* mice (N = 3) and *Naa10*^+/+^ mice (N = 3). Liver and heart lysates were obtained from each mouse for immunoblotting; each liver and heart sample were replicated three times. Blots were stained for total protein post-transfer; after total protein stain removal, blots were incubated with anti-NAA10 MAb and anti-rabbit secondary). **A)** Representative western blot of NAA10 in liver lysates. **B)** Representative western blot of NAA10 in heart lysates. **C)** Quantification of NAA10 signal normalized to total protein. Short black crossbar indicates mean NAA10 signal normalized to total protein (±SD, 2-way ANOVA, F-statistic = 12.52 on 3 and 32 DF, *P < 0.05).

**Fig 2.**
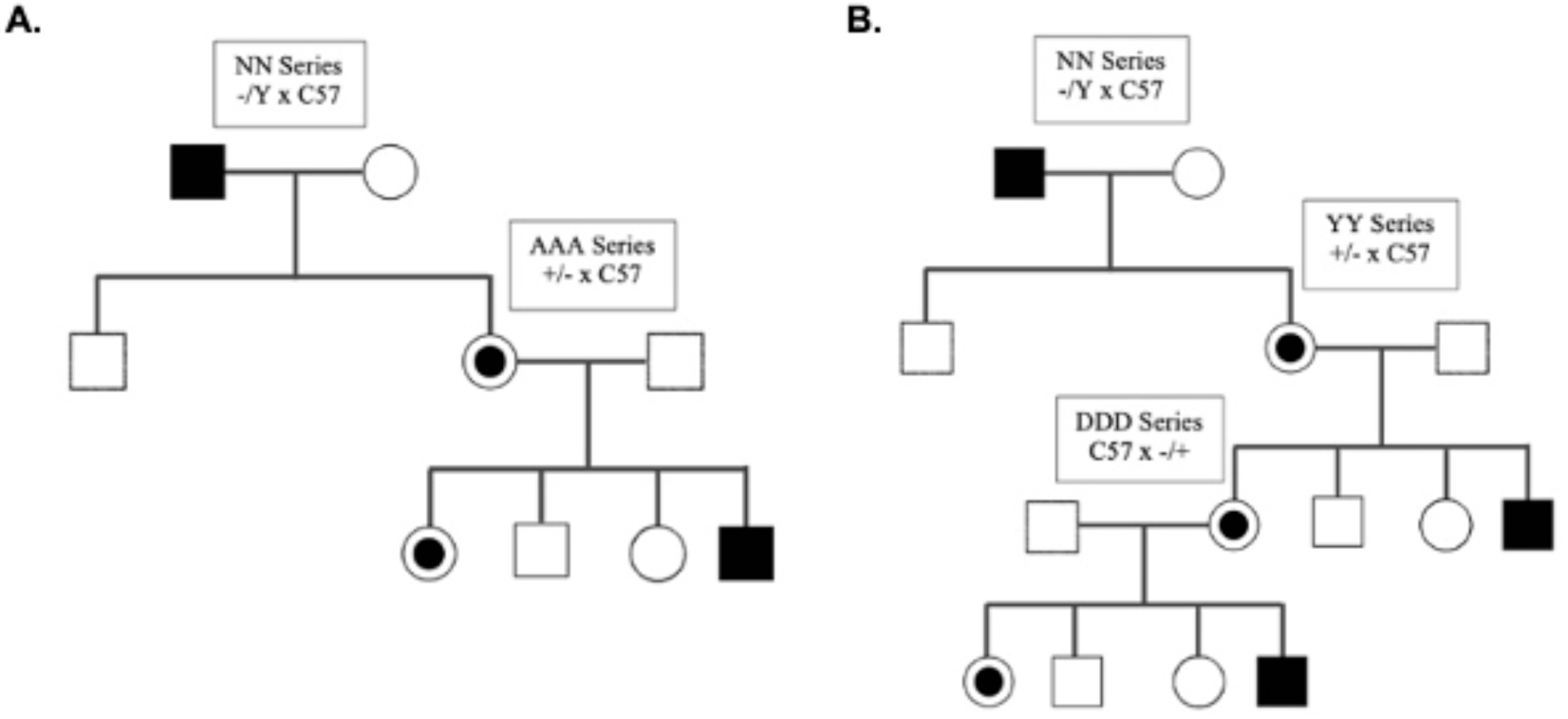
Matings used to generate offspring and embryos. The *Naa10* knockout allele is inherited paternally in the NN series, whereas it is inherited maternally in the YY series. For embryo dissections AAA and DDD series, the allele is maternally inherited for both, but the origin of the allele is from the grandfather for the embryos from the AAA series and from the grandmother for the embryos in the DDD series. These letter combinations (like NN, YY, AAA, DDD) are arbitrary and assigned based on internal lab databases.

The Mendelian ratios for embryos from the AAA and DDD matings with the predicted genotypes are shown in **Table 1**, along with the data combined for AAA and DDD matings. The embryonic age ranged from E9.5 to E16.5, with the bulk of these matings falling between E9.5 to E13.5, as confirmed by Theiler staging comparison. There was some small degree of embryonic lethality (∼20-25%) for *Naa10^−/y^* mice in both mating series, which differs from its absence in a somewhat mixed genetic background of mice bred in an animal facility in Korea, after at least 6 backcrosses to C57BL/6J [21]. Mendelian ratio calculations suggest neonatal lethality exist within both AAA and DDD combined populations where the observed percentage of *Naa10* KO males deviated from the expected 25% to only 16.9% (Table 1).

**Table 1.** Genotypes of E9.5-E13.5 embryos from different mating series.

| ***Naa10<sup>x/-</sup>* female mice crossed to *C57BL/6J* male mice (AAA series)** |  |  |  |  |  |  |
| --- | --- | --- | --- | --- | --- | --- |
| Genotype<br>(Expected Mendelian %) | *Naa10<sup>+y</sup>*<br>(25%) | *Naa10<sup>-y</sup>*<br>(25%) | *Naa10<sup>+/+</sup>*<br>(25%) | *Naa10<sup>-/+</sup>*<br>(25%) | Unable to<br>genotype | Resorptions |
| Embryos + resorptions<br>(n=98) | 19 (19.6%) | 16 (16.5%) | 24 (24.7%) | 28 (28.6%) | 1 (1.0%) | 10 (10.3%) |
| ***Naa10<sup>-x</sup>* female mice crossed to *C57BL/6J* male mice (DDD series)** |  |  |  |  |  |  |
| Genotype<br>(Expected Mendelian %) | *Naa10<sup>+y</sup>*<br>(25%) | *Naa10<sup>-y</sup>*<br>(25%) | *Naa10<sup>+/+</sup>*<br>(25%) | *Naa10<sup>-/+</sup>*<br>(25%) | Unable to<br>genotype | Resorptions |
| Embryos + resorptions<br>(n=75) | 21 (28.00%) | 13 (17.33%) | 17 (22.67%) | 17 (22.67%) | 0 (0%) | 7 (10.29%) |
| **Heterozygous *Naa10<sup>x/-</sup>* or *Naa10<sup>-x</sup>* female mice crossed to *C57BL/6J* male mice (AAA + DDD series)** |  |  |  |  |  |  |
| Genotype<br>(Expected Mendelian %) | *Naa10<sup>+y</sup>*<br>(25%) | *Naa10<sup>-y</sup>*<br>(25%) | *Naa10<sup>+/+</sup>*<br>(25%) | *Naa10<sup>-/+</sup>*<br>(25%) | Unable to<br>genotype | Resorptions |
| Embryos + resorptions<br>(n=173) | 40 (23.3%) | 29 (16.9%) | 41 (23.8%) | 45 (26.2%) | 1 (0.58%) | 17 (11.0%) |

Expected and observed Mendelian ratio of genotypes in offspring from crosses, with paw tattoos and tails taken around day 6 of life.

We next examined possible maternal effect lethality. The *Naa10* allele of our AAA mating series is maternally inherited, with the mother of the dissected embryos having inherited the allele from her father (i.e., the maternal grandfather of the AAA embryos), as displayed in the breeding scheme. The number of our AAA embryos+resorptions (n=98) versus (n=16) is much larger than in the prior study [52] (**Table 1**). There is no statistically significant maternal effect lethality for the heterozygous *Naa10^−/X^* female mice (**Table 2**). **Fig 3a** shows weight gain of dams measured before their mating to their respective sire until the day of necropsy when embryos were harvested, dissected, and measured. *t*-tests comparing the weight gains of C57BL/6J and AAA dams at each length of pregnancy showed no significant statistical difference. **Figs 3B** and **3C** show there is no statistically significant difference in either area or weight of wildtype embryos as compared to *Naa10* knockout embryos in both males and females. Given the larger number of available embryos, within-litter analysis for E12.5 embryos was conducted, and **Figs 4A** and **4B** show four litters only containing E12.5 embryos. The wildtype embryos were greater in area than the knockout mice; however, *t*-tests prove there was no statistical difference between the embryo areas and weights of the wildtype versus knockout embryos. Representative images of the embryos are shown in **Fig 4C**.

**Fig 3.**
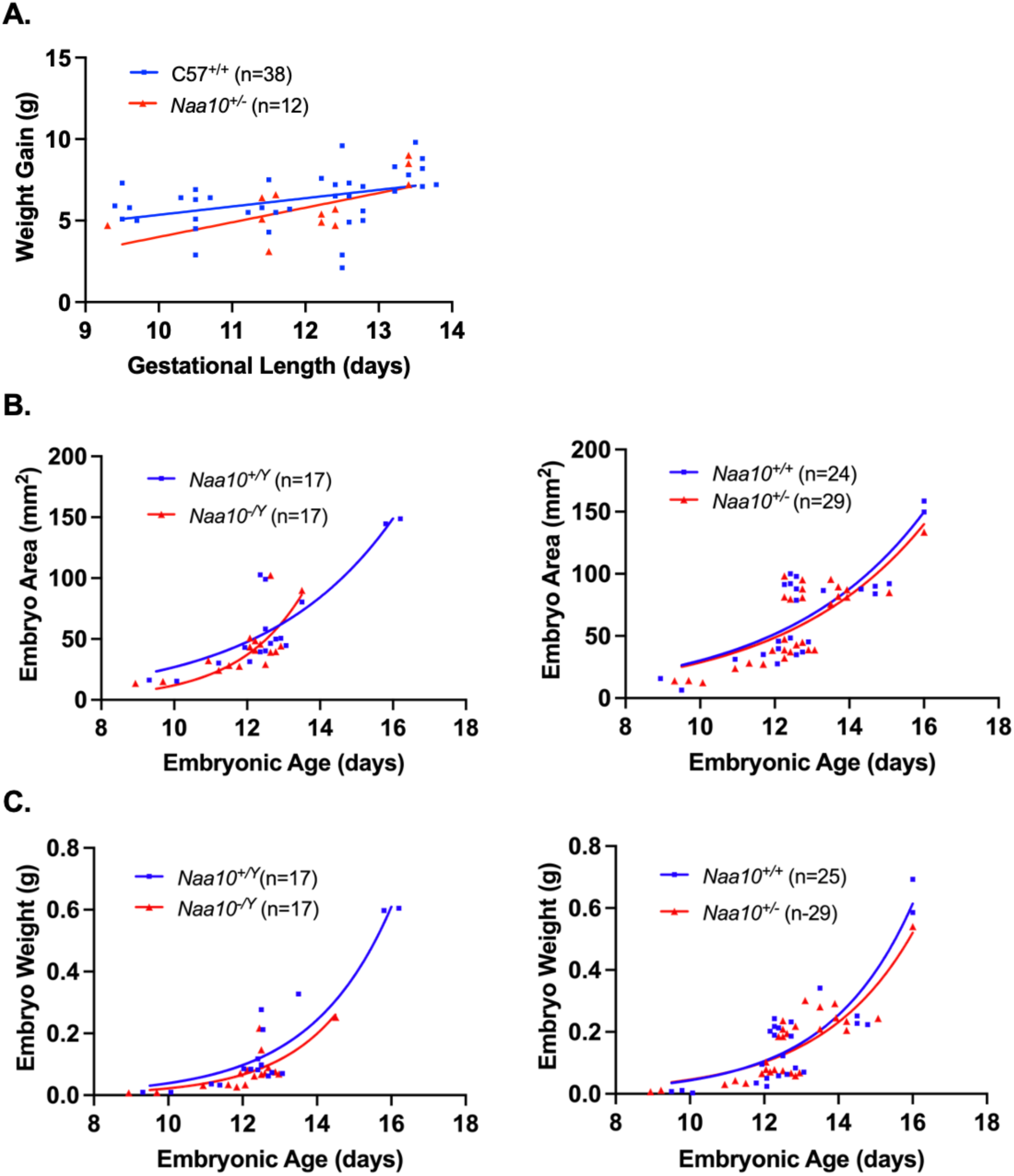
Embryonic phenotypes of *Naa10* knockout mice after 20 backcrosses, maternally inherited but originating from the maternal grandfather, and represented here as AAA. **(A)** *Naa10*^+/−^ and *Naa10*^+/+^ dams were weighed and compared at various lengths of gestation. However, there was no statistically significant difference between both simple linear regression lines that represent the weight gains of *Naa10*^+/−^ and *Naa10*^+/+^ dams at their respective lengths of pregnancy (P>0.05). **(B)** Measured embryo surface areas of male (left) and female (right) versus embryonic age. The areas of *Naa10*^−/Y^ and *Naa10*^+/−^ embryos are not statistically different from those of wildtype (*Naa10*^+/Y^ and *Naa10*^+/+^) embryos (P>0.05). **(C)** Embryo weights of male (left) and female (right) versus embryonic age were determined by measurement. The weights of *Naa10^−/Y^* and *Naa10*^+/−^ embryos are not significantly different from those of wildtype embryos (P>0.05). Graphs **B**-**C** were produced using nonlinear exponential regression modeling.

**Fig 4.**
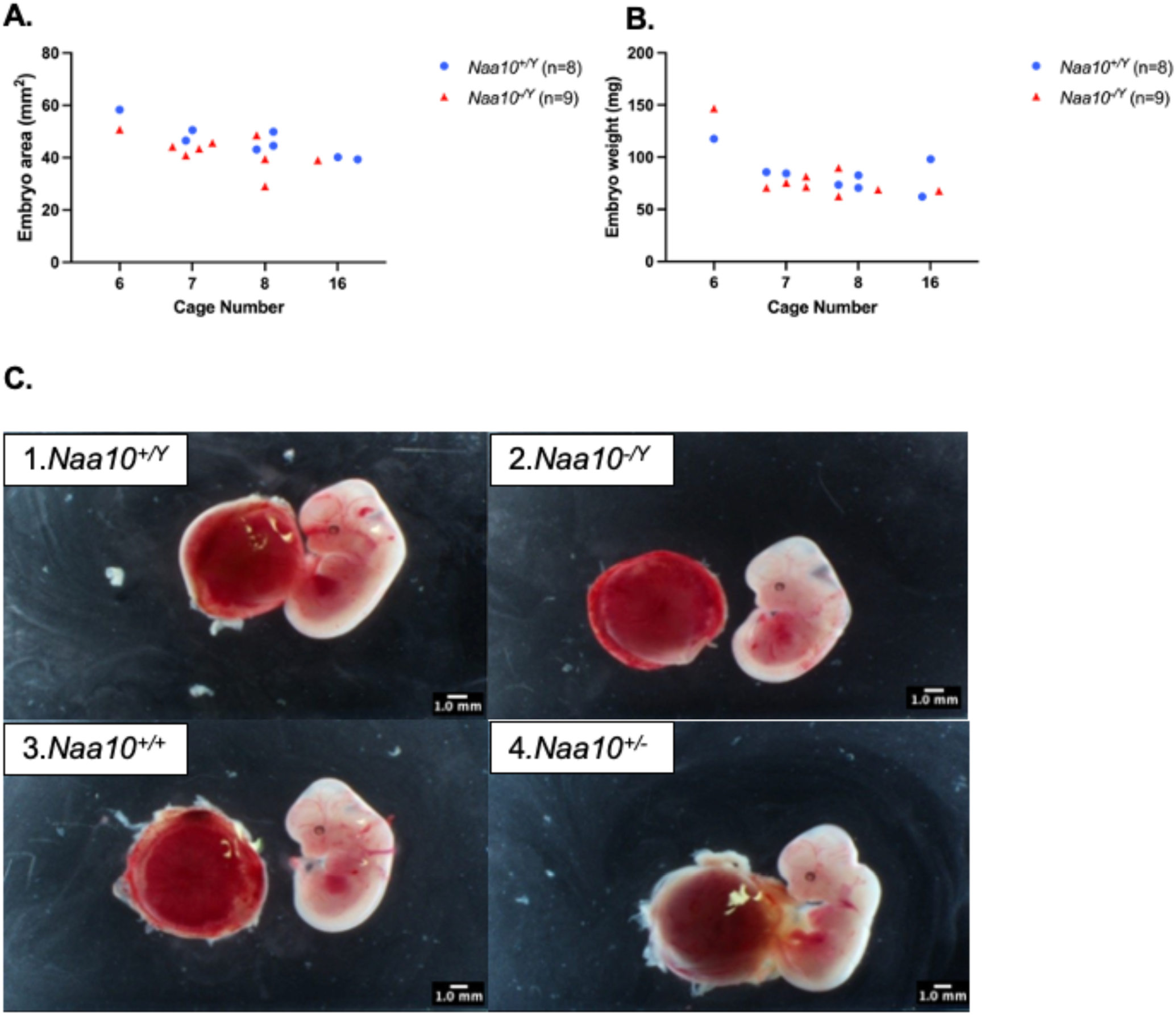
Within-litter analyses for embryos that inherited knockout for *Naa10* maternally and from the maternal grandfather. **(A)** Male embryo areas age E12.5 are graphed and grouped by litter, which were in cage #6, 7, 8 and 16). Two of the four litters contained more than one *Naa10*^−/Y^ embryo. In all four litters, the embryo with the smallest area was the *Naa10*^−/Y^. **(B)** Male embryo weights age E12.5 are graphed and grouped by litter. Two out of four litters contained more than one *Naa10*^−/Y^ embryo. In two out of four litters, the embryo with the lowest weight was the *Naa10*^−/Y^. **(C)** Pictures of E12.5 embryos from litter AAA8: *Naa10*^+/Y^ (1), *Naa10*^−/Y^ (2), *Naa10*^+/+^ (3), *Naa10*^+/−^ (4).

**Table 2.**
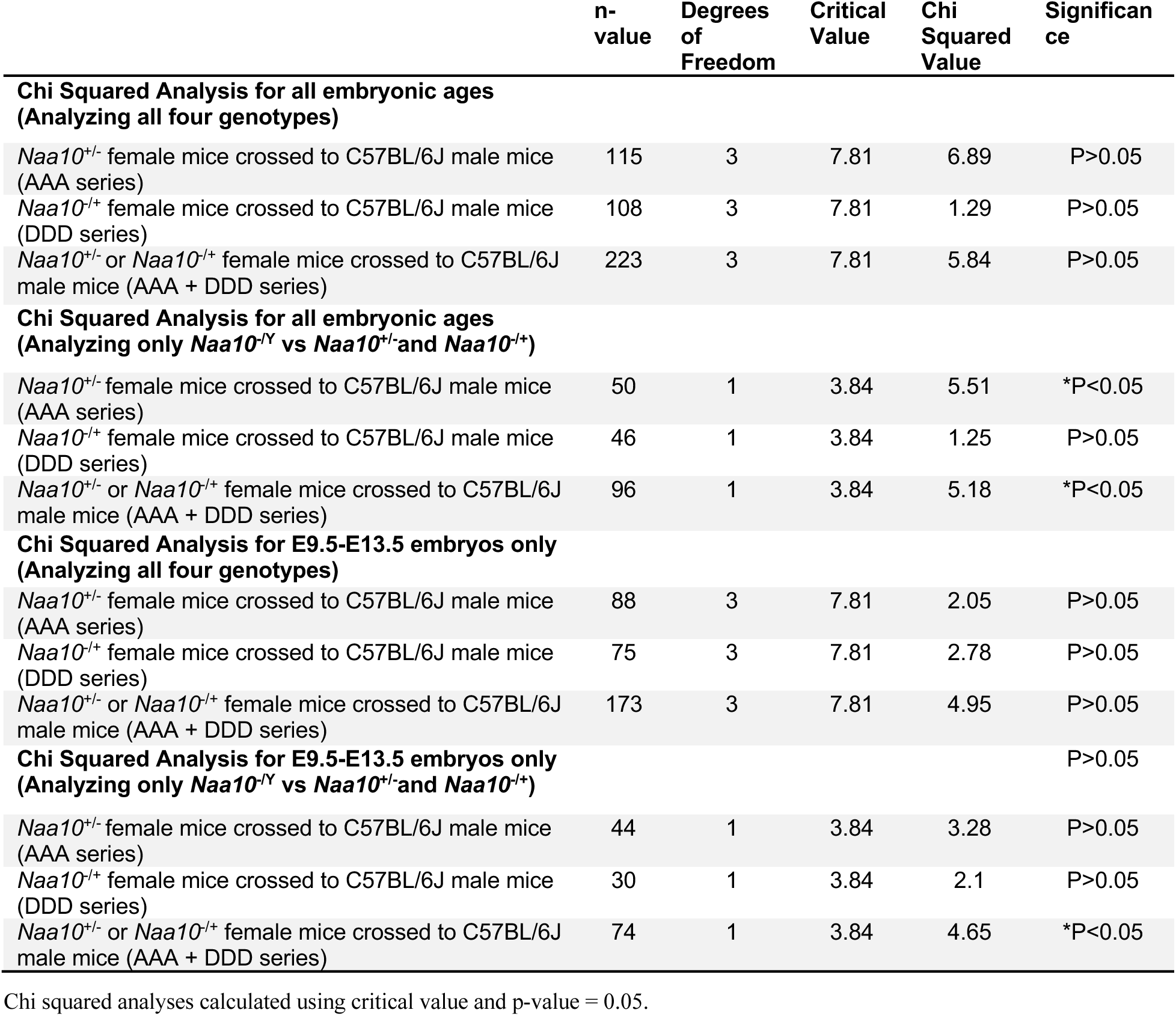
Chi Squared Analyses.

The *Naa10* allele in our DDD mating series is also inherited maternally, but with the mother of these embryos having inherited the allele from her mother, i.e., originating from the maternal grandmother. There is no maternal effect lethality for the heterozygous *Naa10^−/^*^X^ female mice (**Table 1** and **Table 2**). **Fig 5A** shows the weight gain of DDD dams and weight gain of dams from the C57BL/6J line. The weight gain values are measured starting from mating date and day of dissection. *T*-tests comparing the weight gain of dams at each length of pregnancy showed no statistically significant difference (P>0.05). **Figs 5B** and **5C** compare embryo areas and weights of *Naa10* knockout male and female embryos compared to the area and weight values of wildtype embryos. There was no statistically significant difference (P>0.05). Within litter analyses were performed on our DDD embryos where there was no statistical difference caused by variation among different litters. **Fig 6A** shows three litters only containing E12.5 embryos. The smallest area was consistently measured in the *Naa10* knockout embryo. **Fig 6B** shows the weights of the embryos in the three litters. In two of the three litters, the smallest weighing embryo was a *Naa10* knockout. **Fig 6C** shows representative images of the DDD embryos.

**Fig 5.**
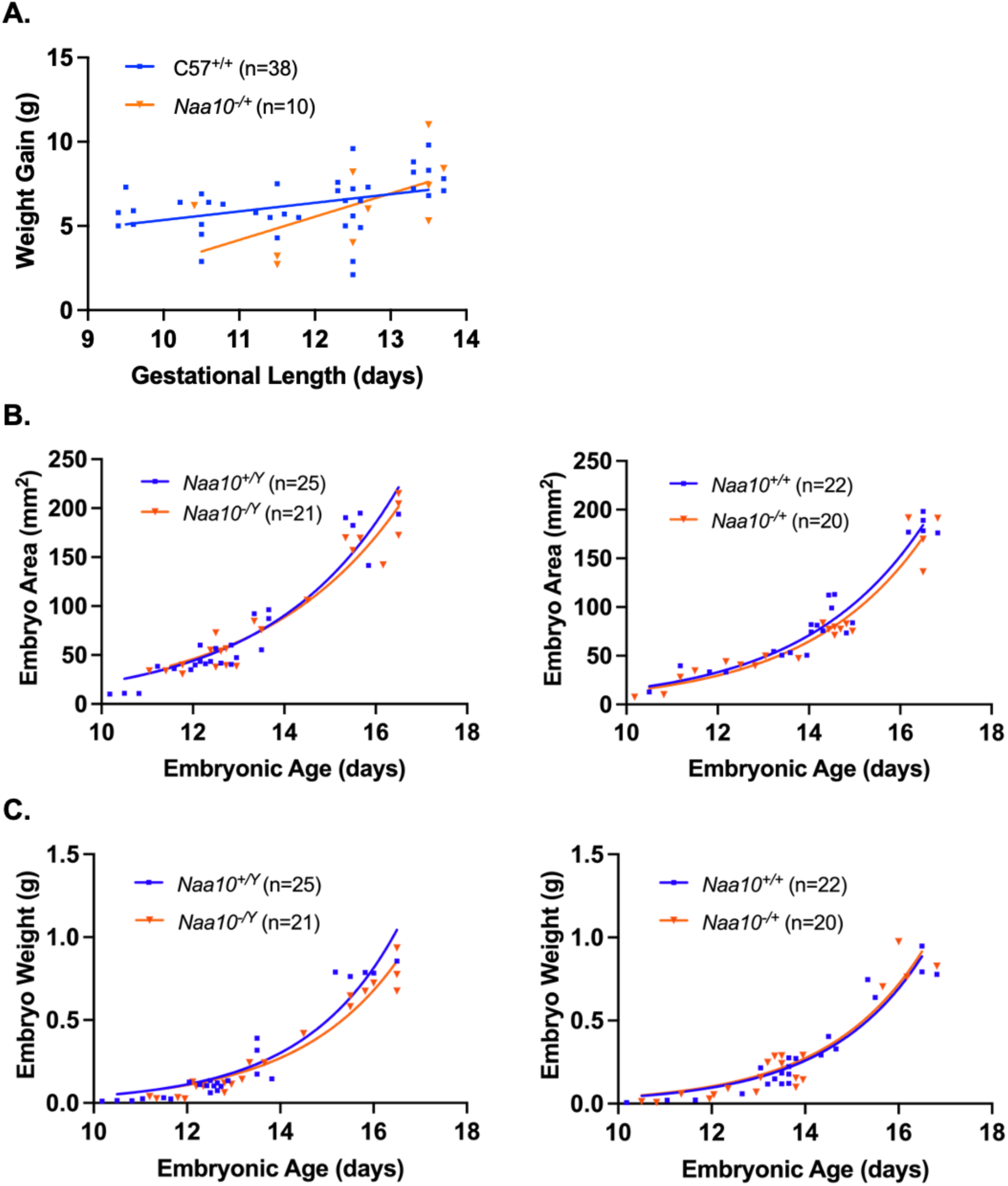
Embryonic phenotypes of *Naa10* knockout mice after 20 backcrosses, maternally inherited but originating from the maternal grandmother, and represented here as DDD. **(A)** *Naa10*^−/+^ and *Naa10*^+/+^ dams were weighed and compared at various lengths of gestation. There was no statistically significant difference between both simple linear regression lines that represent the weight gains of *Naa10*^+/−^ and *Naa10*^+/+^ dams at their respective length of pregnancy. **(B)** Measured embryo surface areas of male (left) and female (right) versus embryonic age. The areas of *Naa10*^−/Y^ and *Naa10*^−/+^ embryos are not significantly different from those of wildtype embryos (P>0.05). **(C)** Embryo weights of male (left) and female (right) versus embryonic age were determined by measurement. The weights of *Naa10*^−/Y^ and *Naa10*^−/+^ embryos are not statistically significantly different from those of wildtype (*Naa10*^+/Y^ and *Naa10*^+/+^) embryos (P>0.05). Graphs **B**-**C** were produced using nonlinear exponential regression modeling.

**Fig 6.**
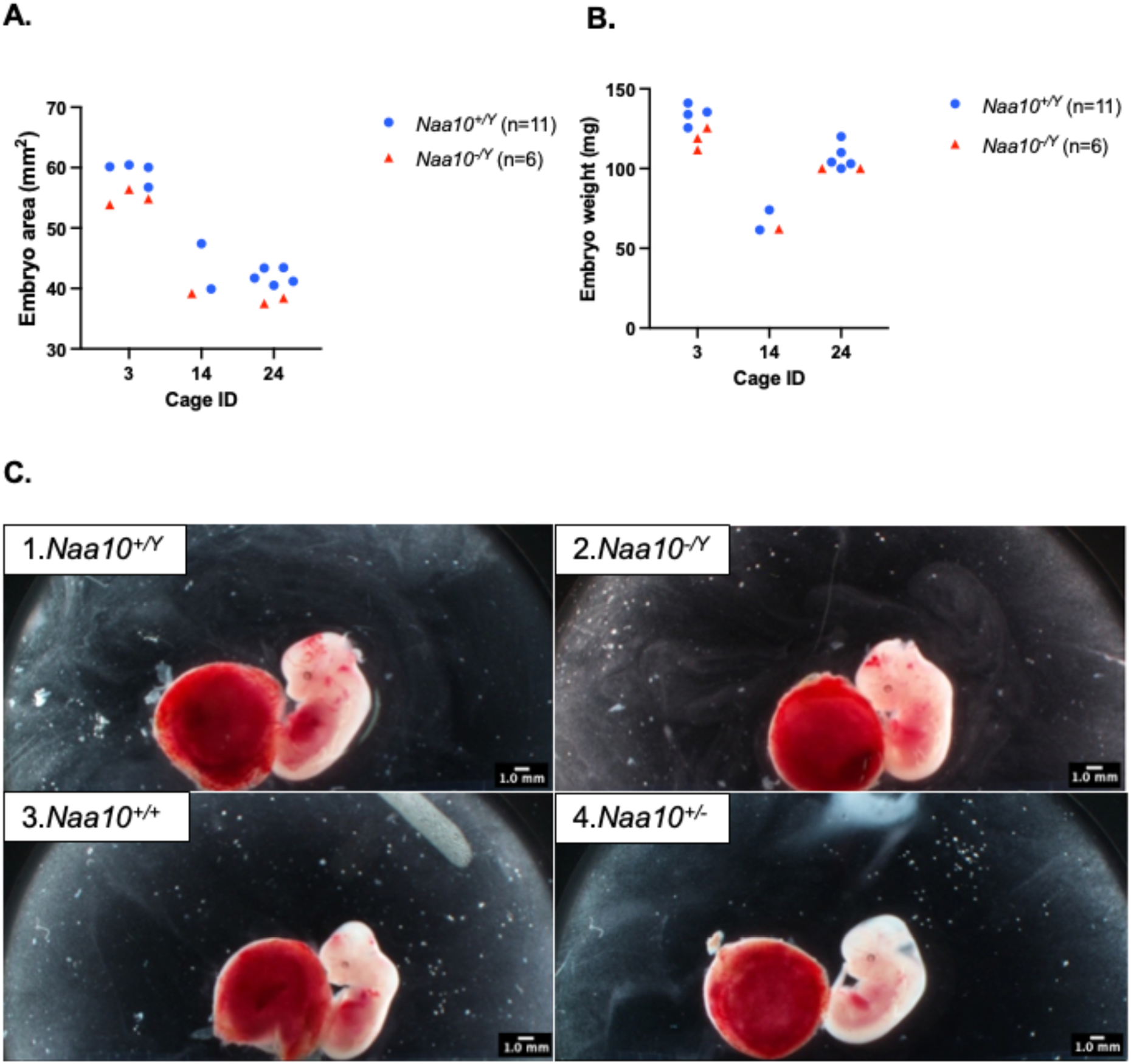
Within-litter analyses for embryos that inherited knockout for *Naa10* maternally and from the maternal grandmother. **(A)** Male embryo areas age E12.5 are graphed and grouped by litter; two of the three litters contained more than one *Naa10*^−/Y^ embryo. In all three litters, the embryo with the smallest area was the *Naa10*^−/Y^. **(B)** Male embryo weights age E12.5 are graphed and grouped by litter. Two out of three litters contained more than one *Naa10*^−/Y^ embryo. In two out of three litters, the embryo with the lowest weight was the *Naa10*^−/Y^. **(C)** Pictures of E12.5 embryos from litter DDD14: *Naa10^+/Y^* (1), *Naa10*^−/Y^ (2), *Naa10*^+/+^ (3), *Naa10*^−/+^ (4).

**Fig 7A** shows no difference in weight gain of the dams’ in both the AAA and DDD series matings. The measured embryo areas were not statistically different when analyzed by genotypes. Finally, data of the two breeding groups ranging from E9.5 to E13.5 were combined and shown at the bottom of **Table 1**. The difference in Mendelian ratios based on the number of heterozygous *Naa10*^−/X^ female mice (n=45) compared to the number of wild type male mice (n=40) is not statistically significant.

**Fig 7.**
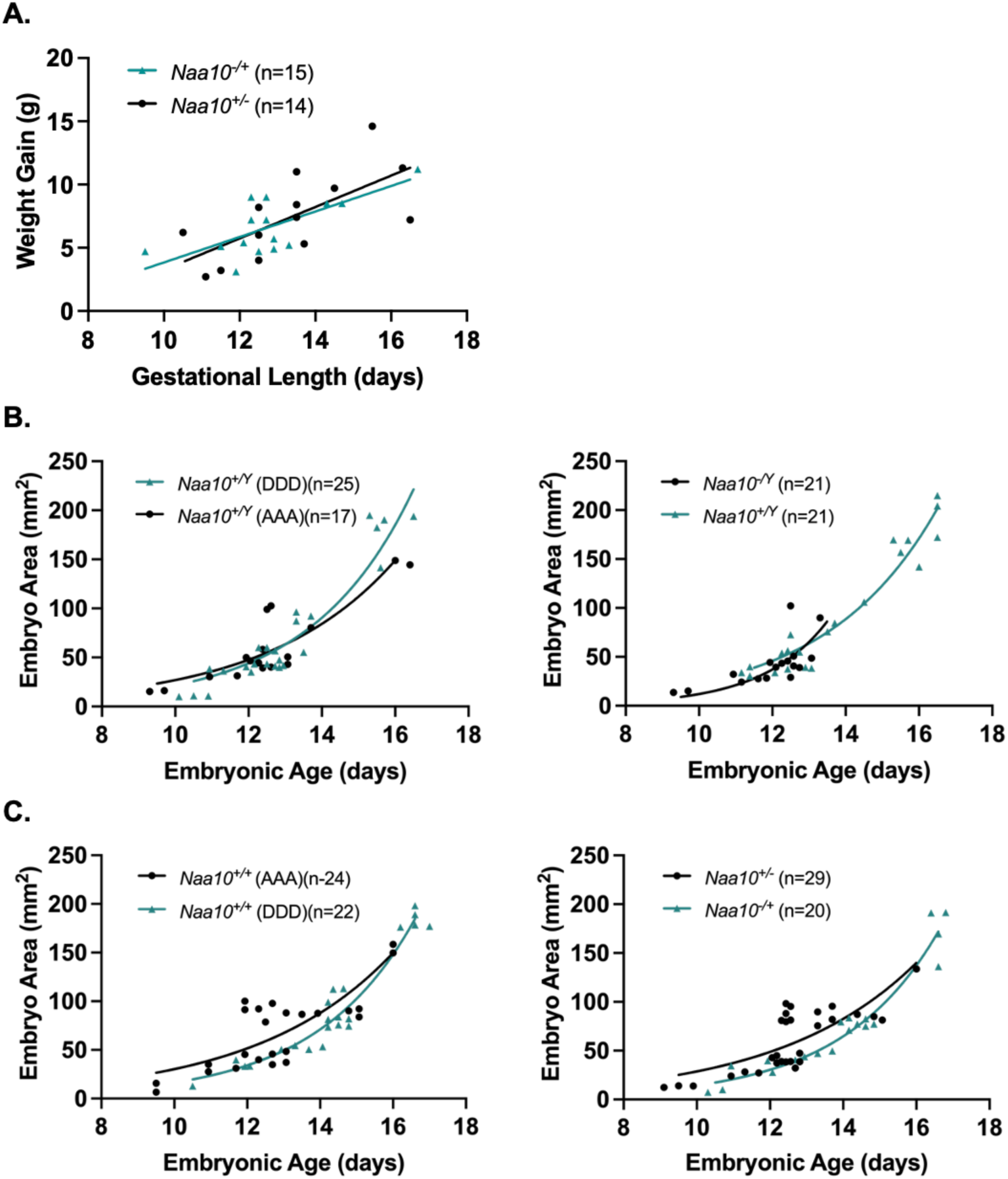
Embryo area versus embryonic age for embryos with maternally inherited *Naa10* knockout allele, where the allele originated from the maternal grandmother (DDD) or the maternal grandfather (AAA). **(A)** Weight gain of *Naa10*^−/+^ (DDD dams) and *Naa10*^+/−^ (AAA dams) at various lengths of gestation were compared. We observed no statistically significant difference between both simple linear regression lines that represent the weight gains of dams from both mating series at each length of pregnancy (P>0.05). **(B)** Left: Embryo area versus embryonic age for DDD vs. AAA *Naa10*^+/Y^ males. Right: Embryo area versus embryonic age for DDD vs. AAA *Naa10*^−/Y^ males. **(C)** Left: Embryo area versus embryonic age for DDD vs. AAA *Naa10*^+/+^ females. Right: Embryo area versus embryonic age for DDD *Naa10^−/+^* vs. AAA *Naa10^+/−^* females. There are no statistically significant differences between the embryo areas for each genotype from each mouse line (P>0.05). Graphs **B**-**C** were produced using nonlinear exponential regression modeling.

### Pleiotropic Effects of *Naa10* Knockout on Postnatal Mice

This current study continues our analysis of the original *Naa10*^−/Y^ knockout strain of live born mice, as previously reported [21], but now genetically inbred with more than 20 backcrosses to C57BL/6J (Jackson Laboratories, Bar Harbour, ME). Surviving *Naa10*^−/Y^ mice showed decreased body weight when compared to *Naa10*^+/Y^ (WT) mice throughout the course of postnatal development. **Fig 8** charts the body weights of male *Naa10*^−/Y^ and *Naa10*^+/Y^ mice from three to 20 weeks. As displayed in the Fig, *Naa10*^−/Y^ mice consistently had reduced body weights compared to *Naa10*^+/Y^ mice throughout the entire 18-week period where data was collected. The mean body weights of *Naa10*^−/Y^ mice presented with lower than the mean body weight when compared to *Naa10*^+/Y^ mice. To further assess whether these differences in body weights were significant, *t*-tests were performed comparing the two groups’ weight data each week. Analysis shows a statistically significant difference (*P<0.05) between the weights of *Naa10*^+/Y^ and *Naa10*^−/Y^ mice in 12 out of 18 weeks. Our observations show that the *Naa10*^−/Y^ mice had significantly less body weight than the *Naa10^+/Y^* mice throughout most of their postnatal development with the exception of six out of the 18 weeks; specifically in weeks five and 16 through 20. During these six weeks there was no statistically significant difference (P>0.05); however, sample sizes were low which may have led to higher variability and lack of significance in these weeks. **Fig 8** shows a phenotypic difference in body weight where the KO mouse is noticeably smaller than the WT littermate mouse. This is similar to the human condition, as some individuals with Ogden syndrome are small in weight and have short stature, whereas a few humans develop at typical size [53].

**Fig 8.**
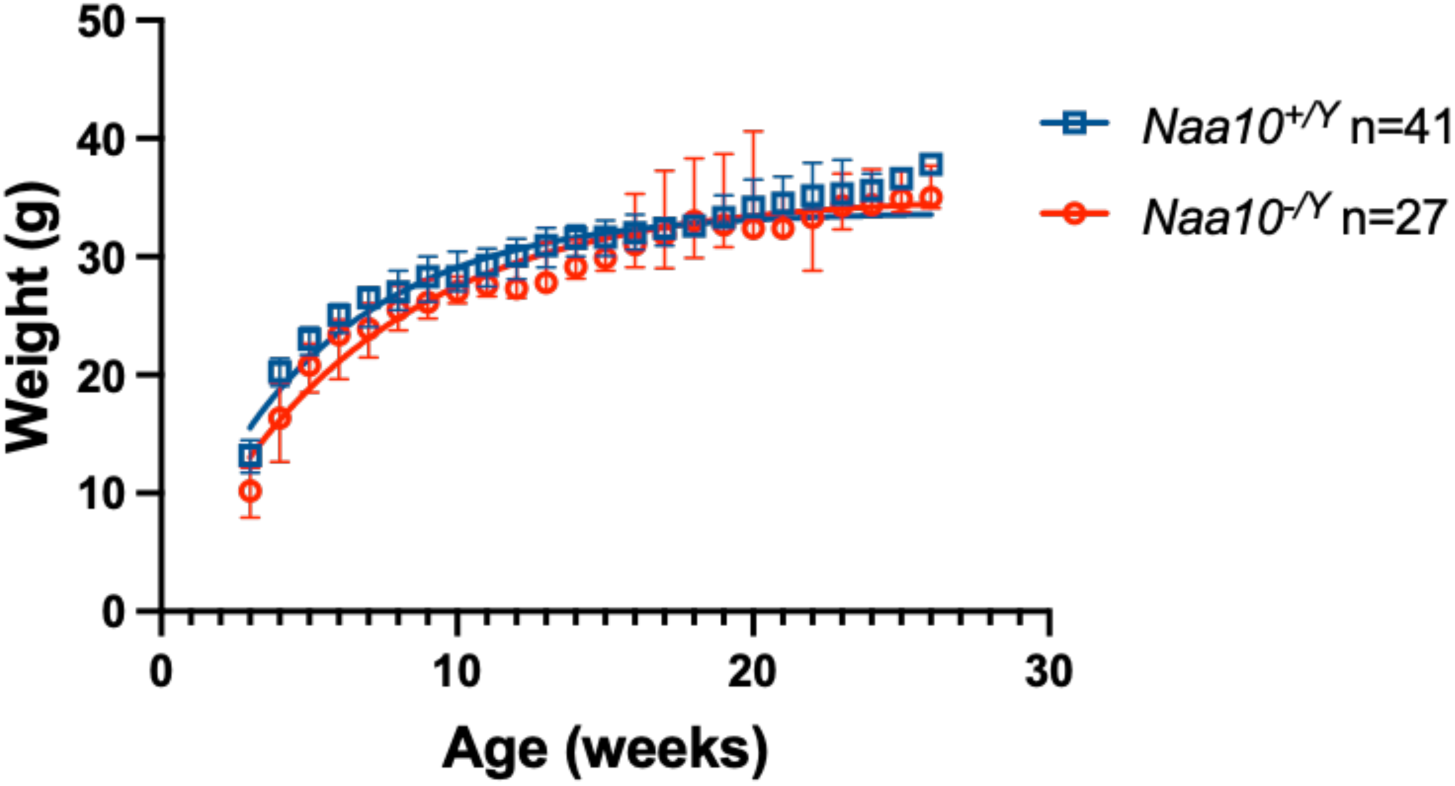
Phenotypic weight data of neonatal *Naa10^−/Y^*(KO) and *Naa10^+/Y^*(WT) mice after 20 backcrosses. Male mice body weights were obtained from 3-20 weeks. Twelve out of 18 weeks showed a statistically significant difference between both nonlinear exponential regressions of body weights when comparing KO mice to wildtype littermates (*P<0.05).

Other phenotypic defects present in the *Naa10* knockout mice include hydrocephalus, piebaldism, and hydronephrosis (**Table 3**), which are consistent with previous findings [21].

**Table 3.** Percentage of phenotypes observed in *Naa10^−/Y^* mice vs. C57BL/6J male mice.

| Phenotypes | Piebaldism | Hydrocephalus | Hydronephrosis | White digits and tail tips. |
| --- | --- | --- | --- | --- |
| Percentage in *Naa10<sup>-Y</sup>* mice | 97% (28/29) | 17% (5/29) | 27% (3/11) | 100% (29/29) |
| Percentage in male C57BL/6J mice | 0% | 1.6% (3/191) | Never observed in routine necropsies | 0% |

**S12A Fig** shows an *Naa10^−/Y^* mouse (left) with hydrocephaly as compared to a wildtype littermate with a normally developed skull as well as normal body composition. Hydrocephalus was observed in 17% (5/29) of *Naa10^−/Y^* mice. **S12B Fig** displays piebaldism on the ventral abdomen of a *Naa10*^−/Y^ mouse, which is a consistent phenotype seen in many *Naa10*^−/Y^ mice. This phenotype was observed in 97% (28/29) of *Naa10*^−/Y^ mice. The prior paper reported piebaldism in 100% of *Naa10*^−/Y^ mice [21]. We believe that the one mouse we observed to lack piebaldism may have lost the hypopigmentation during maturation (discussed further below). **S12C Fig** displays hydronephrosis in a *Naa10^−/Y^* mouse. The left kidney is noticeably enlarged, with the left kidney weighing 13.02 grams compared to the right kidney weighing 0.34 grams. **S12C Fig** also shows that the *Naa10*^−/Y^ mouse presenting with hydronephrosis also had an enlarged bladder. The hydronephrosis phenotype was observed in 27% (3/11) of surviving *Naa10*^−/Y^ mice.

White digits and tails can be seen in the *Naa10*^−/Y^ mice shown in **S12A Fig (left)** and **S12B (left)**, as the *Naa10*^−/Y^ mouse exhibits ventral abdominal piebaldism, white tail tips and digits, while its wildtype littermate does not. We observed this phenotype in 100% (29/29) of *Naa10* knockout mice on this pure genetic background, displaying its complete penetrance on that background.

## Discussion

*NAA10*, which encodes the catalytic subunit of the principal NAT complex (NatA) and is mutated in humans affected by OS, is likely an essential gene for eukaryotic development. Initially reported in yeast, other Naa10 (also known as Ard1) orthologues were later identified in humans and various model organisms, including *D. melanogaster, D. rerio, A. thaliana*, and *M. musculus* [55,56]. In *D. melanogaster*, ovary-specific loss of Naa10 causes defective oogenesis in *vnc* mutants. The loss of Naa10 also contributes to cell growth and proliferation failure in wing discs [57]. Morpholino-mediated Naa10 knockdown in *D. rerio* increases developmental lethality; further, surviving zebrafish exhibit growth failure and other developmental abnormalities (e.g., bent axes, missing eyes, and missing tails) [58]. In addition to the apparent conservation of Naa10, the NTA profile of plants and animals are broadly similar which supports the necessity of this protein modification in eukaryotes [59,60]. *Naa10* knockout mouse lines also support the necessity of NatA-catalyzed NTA in mammalian development [21,61,62]. Previously, we showed that Naa12 complexes with Naa15 to form functional NatA in Naa10^−/Y^ animals, which explains why these mice were born and were not completely embryonic lethal, at least in the mixed genetic background reported in that paper [21]. Confirming this, double mutants for Naa10 and Naa12 knockout mice were embyronic lethal [21].

The impact of genetic background is supported by our observation that additional null alleles (*Naa10* Δ668 and *Naa10* Δ668-674) on mixed genetic backgrounds (∼ 75% C57BL/6J, 25% DBA/2J (D2)) have far less penetrance for piebaldism and much less (if any) neonatal lethality, whereas the piebaldism was fully penetrant after eight backcrosses to C57BL/6J for the previously published allele [21], along with the previously reported partial penetrance for ventricular septal defects (VSD), atrial septal defects (ASD), persistent truncus arteriosus (PTA) or double outlet right ventricle (DORV) of the mice who died in the first three days of life (n= 6/28, or 21%) [21]. We also previously reported that there was no embryonic lethality for the *Naa10*^−/y^ mice after eight backcrosses to C57BL/6J, except for when the compensating enzyme Naa12 was knocked out [21]. The present study examined this on an inbred C57BL/6J genetic background (20 backcrosses) and in the new environment of the animal facility at IBR, where we did observe some embryonic lethality (∼20-25% of *Naa10*^−/Y^ male mice), even in the presence of *Naa12*. Phenotypic variation due to genetic background has been observed in several genetically engineered mouse models [63]. Alternatively, the transfer of all mice to a new animal colony and environment in March 2019 may have contributed to this embryonic lethality potentially through the effects of maternal stress on *in utero* development [64] or immune system dysfunction [65]. In support of this latter hypothesis, there was a parallel increase in neonatal lethality in the new colony observed for all mice, including the wild type *C57BL/6J* mice (unpublished data). Lastly, the gene targeting scheme presents the possibility that insertion of the neomycin cassette utilized to generate this strain of Naa10 KO mice may have resulted in dysregulation of off-target gene expression - a phenomenon documented in previously published literature [66–71], which could itself be modified by genetic background and environment, although this is not something that we have investigated for our mice.

We have previously noted that *Naa10^−/−^* mutants somewhat phenocopy *Pax3* mutants [21]. *Pax3* knockout mice, which model symptoms of Waardenburg syndrome, lack a transcription factor necessary for cardiac neural crest cell (cNCC) migration and proliferation in the pharyngeal arches and embryonic outflow tract. Without Pax3, inadequate NCC migration results in congenital heart defects attributable to improper remodeling of the arches and outflow tract [72–75]. *Pax3^+/−^* adults have 100% piebaldism, along with variable penetrance for neural crest (NC)-related persistent truncus arteriosus (PTA) or double outlet right ventricle (DORV) with concomitant ventricular septal defects VSDs [72,73,76,77], congenital hydrocephalus [78], and/or skeletal defects due to abnormal somite morphogenesis [79,80]. Of most relevance here, the phenotype of *Pax3* nulls can be modulated by genetic background, in which it has been shown that those mice exhibit 100% mid-gestational lethality due to cardiac neural crest-related deficiencies on C57Bl/6J inbred genetic background [77]. As such, there is modulation of these neural crest-related phenotypes by genetic background. There is a long and very extensive literature regarding the modulation of phenotype by genetic background, some of which is summarized in a book chapter written by one of us [81,82].

A prior study [52] found embryonic lethality in mice on a somewhat mixed genetic background, as that strain was used after “at least six generations of backcross with C57BL/6 mice”, which was noted by the authors to be the sub-strain C57BL/6JNarl, first established at the Animal Center of National Research Institute from the Jackson Laboratories in 1995. However, as noted, the different environments of the animal facilities may play some role; therefore, embryonic lethality may be somehow more apparent even when the mice are not fully inbred. Also, the previous study [52] used the Cre/loxP system to generate the *Naa10* KO mice, where a floxed *Naa10* female mouse was crossed with the Ella-Cre transgenic male mouse expressing Cre recombinase for germ line deletion of loxP-flanked *Naa10*, whereas the Naa10^−/y^ mice were made using standard gene-targeting methods without the use of Cre recombinase [21,54]. It is not clear how Cre recombinase could impact the phenotype in the strain maintained at the Taiwan facility. It is notable that the other group did not comment on whether their mice have piebaldism or any skeletal defects, so it is not clear if the mice possess these phenotypes [52,62].

We report that we cannot replicate the previously reported “maternal effect lethality” and we cannot find any correlation between the sizes of *Naa10*-null embryos and placental weight [52]. It is possible that additional study of earlier embryonic ages might reveal whether the embryonic lethality starts internally within the embryo or if it is mediated instead by some placental abnormality, as previously claimed [52]. Since this prior publication, there have been no other publications replicating these effects or the genomic imprinting findings of the paper. There is also still no crystal structure or any other structure showing that *Naa10* has any direct DNA-binding domain or activity. The authors speculated in their paper that the various phenotypes of the mice (growth retardation, embryonic lethality, brain disorder, and maternal-effect lethality) might be caused by a previously unappreciated role for *Naa10* in DNA methylation and genomic imprinting. Notably, that study was published prior to the discovery of *Naa12* [21], and those results should be re-evaluated in the context of this compensating enzyme. The prior study utilized a small number of mice and samples for the various phenotypes in the mice such as “maternal effect lethality” and were able to only use two embryos of each genotype for the imprinting analyses [52]. It thus remains an open question in the field whether the various replicated phenotype findings in the mice (such as variable amounts of piebaldism, skeletal defects, small size, hydrocephaly, and hydronephrosis) are due to decreased amino-terminal acetylation of certain key proteins or whether some effect related to DNA methylation or genomic imprinting plays a role. Other groups have published that NAA10 might acetylate lysine side chains as a lysine acetyltransferase (KAT), although this is controversial [83]. Future studies are needed to identify acetylated proteins and to study their mechanism of action in both mouse models and humans.

## Material and Methods

### Experimental animals

All experiments were performed in accordance with guidelines of International Animal Care and Use Committees (IACUC) of Cold Spring Harbor Laboratory (CSHL) and New York State Institute for Basic Research in Developmental Disabilities (IBR).Mice suffering from hydrocephalous or serve cases of ulcerative dermatitis, malocclusion, and microphthalmia due to genetics of the strain humane endpoint intervention was used. Humane endpoint intervention consisted of exposure to CO2 to render the animal unconscious, followed by cervical dislocation to ensure rapid passing. Potential breeding complications such as dystocia and severe fight wounds, partially cannibalized neonates, trampled neonates, loss of body condition, reproductive tumors and other causes of neonatal death, injured or traumatized animals are also sought humane endpoint intervention.

Mice were moved to the IBR facility in March 2019, after being housed at CSHL from 2015-2019. While at CSHL, they were housed as breeding pairs or were weaned and housed by sex in individually ventilated autoclaved caging (Ancare Ventilated Caging, Bellmore, NY). At IBR, animals are maintained in autoclaved cages and bedding with 1/8-inch corn cob bedding (The Andersons, Maumee, OH) and were fed a closed-formula, natural-ingredient, γ-irradiated diet (PicoLab Mouse Diet 5058, Purina LabDiet, St. Louis MO) *ad libitum*, and received tap municipal water in polysulfone bottles (Thoren). Mice receive a sterile supplement Love Mash Rodent Reproductive Diet (Bio-Serv; Flemington, NJ). A complete cage change was performed every 7-10 days within horizontal laminar flow cage change station (model Nu602-400Class II TypeNuaire, Plymouth, MN). The room was maintained on a 10:14-h light: dark cycle with a relative humidity of 30 – 70%, and room temperature ranging from 69-78°F. At IBR, since March 2019, mice were housed in a specific pathogen free room of a conventional animal facility in accordance with the *Guide for the Care and Use of Laboratory Animals* (8th edition)[84]. Mice were housed as breeding pairs or were weaned and housed by sex in individually ventilated autoclaved caging (no. 5, Thoren Caging Systems, Hazelton, PA).

Rodent health monitoring assessment is performed three times a year. Mice in this colony are specific pathogen free for the following; astroviruses types 1 and 2, new world Hantaviruses, lymphocytic choriomeningitis virus, mouse adenovirus types 1 and 2, mouse hepatitis virus, ectromelia virus, mouse kidney parvovirus (murine chapparvovirus), mouse parvovirus, minute virus of mice, mouse rotavirus (epizootic diarrhea of infant mice virus), pneumonia virus of mice, reovirus, Sendai virus, Theiler meningoencephalitis virus; beta Strep Groups A, B, C and G, *Bordetella bronchiseptica*, *Bordetella pseudohinzii*, *Campylobacter* spp., ciliary-associated respiratory bacillus (*F. rodentium*),*Corynbacterium kutscheri*, *Klebsiella oxytoca*, *Klebsiella pneumoniae*, *Mycoplasma pulmonis*, *Citrobacter rodentium*, *Rodentibacter* spp., *Chlamydia muridarum, Pseudomonas aeruginosa*, *Salmonella* spp., *Streptobacillus moniliformis*, *Proteus mirabilis*, and *Clostridium piliforme*; *Pneumocystis*, *Giardia* spp. and *Spironucleus muris*; and fur mites (*Myobia musculi*, *Myocoptes musculinis*, and *Radfordia affinis*) and pinworms (*Syphacia* spp. and *Aspiculuris* spp.) all of which is based on multiplex polymerase chain reaction pooled from oral swabs, pelt swabs, and fecal pellets obtained directly (representing 10-15% of cages) and serologic panels from a subset (representing ∼2% of cages). PCR and serology were performed by Charles River Laboratories (Wilmington, MA).

### Generation and genotyping of *Naa10* knockout mice

The *Naa10* knockout (KO) mice were generated as previously described (Yoon, H., et al). The progeny was backcrossed to C57BL/6J for more than 20 generations. This was confirmed with genome scanning at the Jackson Laboratory. The stock of C57BL/6J was replenished annually from Jackson Laboratory (JAX) to avoid genetic drift from the JAX inbred line. Paw tattoo and tail genotyping was performed on day 5 or 6 of life, so as not disturb the litters and thus not increase the risk for maternal rejection of the litter. The primers used for *Naa10 KO* and *Naa10^tm1a^* genotyping were Naa10-F: 5’-cctcacgtaatgctctgcaa-3’, Naa10-neo-F: 5’-acgcgtcaccttaat-atgcg-3’, Naa10-R: 5’-tgaaagttgagggtgttgga-3’ (**S8 Table)**.

### Generation and genotyping of *Naa10* minigene mice

Standard methods were used to generate and select ES clones that were used for blastocyst microinjection and generation of chimeric mice. Chimeric mice were mated with C57BL/6 mice, and germ-line transmission of targeted alleles was detected by PCR. Primers used were: MG1F: 5’-GTCGACGGCTCAGCATGAAGA; Lox1: 5’-AGCTCCTATCGTCCTTTCCCTGC; SQ2: 5’-AACTATGGCCAGCTTGCTATG; and PT4: 5’-TCTCCAGTCTACCTCTACCAAACCC. Ser37Pro Genotyping PCR: MG1F-PT4, 980 bp product; Ser37Pro mutation activation PCR: Lox1-MG1F (730 bp) or Lox1-SQ2 (1150 bp).

### Generation of *Naa10* (Gm16286, UniProt: Q9CQX6) indel knockout mice

The mice were made using standard methods by microinjection of CRISPR reagent mix into zygotes obtained from the mating of B6D2F1 females (i.e., 50% C57BL/6J, 50% DBA/2J (D2)) females to inbred C57BL/6J males. The guide RNA was produced and validated from Horizon using a Cel1-nuclease assay, and the most active guide was selected, including the targeting cr-RNA sequence and the tracrRNA portion. The indels were transmitted by breeding again to inbred C57BL/6J males, and the resulting progeny were interbred on a mixed genetic background of approximately 12.5% DBA/2J (D2) / 87.5% C57BL/6J, for use in the reported experiments. Genomic DNA was isolated from paw and tail. DNA was screened for mutations using PCR and Surveyor assay [85], followed by Sanger sequencing of selected clones and the use of CRISP-ID [86] to identify putative deletions.

### Quantitative PCR

Organs from mice were dissected >2 months after birth. 70-120 mg tissue (heart/kidney/liver) were lysed in 5 μl/mg tissue RIPA buffer (Sigma) with 1x Complete protease inhibitors and 1 U/μl Superase In (Thermo Scientific, Waltham, MA, USA) using Fisherbrand Pellet Pestle Cordless Motor. After homogenization debris was removed by centrifugation at 20.800 g for 10 min at 4°C. Protein concentration was determined using APA assay (Cytoskeleton Inc. Denver, CO, USA) and 50 μg total protein were separated on SDS-PAGE followed by western blot. Membranes were stained with anti-Naa10, anti-Naa15 and anti-GAPDH antibodies (all Abcam, Waltham, MA,USA). For RNA purification, 30 μl clarified lysate were mixed with 70 μl RNase free water and RNA isolated using the RNeasy Mini Kit (Qiagen, Germantown, MA, USA) according to the manufacturer’s recommendations, including on-column Dnase digest. 1 μg RNA was reverse transcribed using the TaqMan Reverse transcription kit and gene level detection performed using SYBR Green Master Mix (all Thermo Scientific). Relative expression was normalized to GAPDH and ACTB. The following primer pairs were used:

Naa10 (exon2-4) CTCTTGGCCCCAGCTTTCTT & TCGTCTGGGTCCTCTTCCAT

Naa11 ACCCCACAAGCAAAGACAGTG & AGCGATGCTCAGGAAATGCTCT

GAPDH AGGTCGGTGTGAACGGATTTG & TGTAGACCATGTAGTTGAGGTCA

ACTB GGCTGTATTCCCCTCCATCG & CCAGTTGGTAACAATGCCATGT

### Isolation and imaging of mouse embryos

Timed matings were performed by counting the number of days since the male and female were paired. The male mice were left in the cage for three days, prior to removal, giving a three-day window for embryogenesis. Theiler staging was performed for a precise gestational age. Pregnant mice were euthanized at several time points after conception. The embryos were isolated on ice, then washed three times in cold 1% PBS. Embryos were imaged using an Olympus SZX10 with Olympus CellSens imaging software (Center Valley, PA, USA). Both embryos and placentas were measured, weighed, then stored in 10% formalin buffered saline and then stored at four degrees Celsius. Scale bars were created for adult mouse and embryo dissections using software program Fiji by ImageJ (National Institute of Health, Public Domain, BSD-2)

### Whole body CT scanning

CT scans were acquired on a Nanoscan PET/CT scanner at CSHL from Mediso using Nucline v2.01 software. All mice were kept sedated under isoflurane anesthesia for the duration of the scan. Scans were acquired with an X-ray tube energy and current of 70kVp and 280uA respectively. 720 projections were acquired per rotation, for 3 rotations, with a scan time of approximately 11 minutes, followed by reconstruction with a RamLak filter and voxel size 40×40×122µm. The relative mean bone density of the femur from these mice was measured in Hounsfield units using VivoQuant software (v2.50patch2). Briefly, the femur was accurately segmented from the image by first applying a ROI about the bone. Global thresholding, with a minimum of 1000 HU and a maximum of 8000 HU was then applied to accurately segment the femur from the initial ROI. For *ex vivo* analyses, mouse heads were fixed in 10% formalin buffered saline, followed by scanning and reconstruction with 1440 projections per revolution. Cranial volume was measured using VivoQuant software (v2.50patch2), using the spline tool to manually draw around the circumference of the cranium on multiple stepwise 2D slices.

### Western blot

Adult mice were euthanized in a CO_2_ chamber, followed by cervical dislocation. Tissue was dissected and washed in 1% PBS before immediate processing or flash freezing in liquid nitrogen. Fresh or thawed tissue was lysed in RIPA buffer supplemented with protease inhibitor (#R0278, Sigma-Aldrich; #11836170001, Roche) and disrupted using a handheld cordless motorized microtube pestle. Lysate was cleared via centrifugation. Protein quantification was determined via Bradford assay using PrecisionRed Advanced Protein Assay reagent (#ADV01, Cytoskeleton, Denver, CO, USA). Lysate concentration was normalized before dilution in 2X Laemmli Sample buffer with 10% v/v 2-mercaptoethanol. Accordingly, reducing and denaturing SDS-PAGE was conducted on 10% resolving gel (#4561033, Bio-Rad, Hercules, CA, USA) in the Mini-PROTEAN Tetra Cell system. Resolved proteins were transferred to 0.2 µm nitrocellulose membrane (Amersham, Buckinghamshire, UK) using Towbin’s transfer buffer (100 V, 30 min). Membranes were dried at least 15 minutes before reactivating in TBS. Reactivated membranes were stained for total protein using REVERT 700 Total Protein stain kit and scanned wet on Odyssey Classic in the 700nm channel (LI-COR, Lincoln, NB, USA). **S9 Table**.

After scanning, membranes were blocked in 5% non-fat dry milk (1 hr, RT). Blocked membranes were incubated with 1/500 anti-NAA10 Mab (#13357, Cell Signaling Technology, Danvas, MA, USA) diluted in blocking buffer supplemented with 0.1% Tween-20 (overnight, 4°C). After primary incubation, membranes were placed in TBS-T for 5 minutes (3 repetitions). Membranes were then incubated in 1/20,000 goat-anti-rabbit IR800 CW secondary antibody diluted in blocking buffer (1 hr, RT). Stained membranes were washed in TBS-T for 5 minutes (3 repetitions) and rinsed in TBS. Membranes were dried before scanning on the Odyssey Classic. NAA10 signal was normalized to total protein as indicated by REVERT 700 stains; normalized NAA10 signal was quantified in Empiria Studio (LI-COR). Hypothesis testing was conducted in RStudio. Western blot datasets and analyses are available on Github (https://github.com/ajgarcuny).

For Western blotting of the minigene and inverted Naa10 mice, 50-180 mg tissue (liver or heart) was lysed in RIPA buffer (Sigma) with Complete protease inhibitors (5 ul lysis buffer per mg tissue) using Fisherbrand Pellet Pestle Cordless Motor. After homogenization debris was removed by centrifugation at 20,800 g for 10 min at 4°C. Protein concentration was determined using APA assay (Cytoskeleton Inc.) and 50 - 100 μg total protein was separated on SDS-PAGE followed by western blot. Membranes were stained with anti-Naa10, anti-Naa15 and anti-GAPDH antibodies (all Abcam).

For Western blotting on the indel mice, three mice of each genotype (NAA10 wild type [wt], Δ668, or Δ668-674), tissue lysates were prepared from 30-80 μg of liver by mechanical lysis in RIPA buffer. Protein concentration was determined using a commercial version of the Bradford assay, and then normalized to 5 mg/ml in Laemmli SDS-PAGE Sample Buffer. Twenty microliters (i.e., 100 μg) was then loaded on each of two 10% SDS-PAGE gels. After running, gels were transferred to PVDF membranes and then blocked. After blocking, one membrane was incubated with rabbit anti-Naa10 (ab155687, Abcam, Walthan, USA) followed by goat anti-rabbit IRDye 680RD (926-68071, LI-COR, Lincoln, NB, USA), while the other was incubated with mouse anti-GAPDH (Abcam ab9484) followed by goat anti-mouse IRDye 800CW (LI-COR 926-32210). The membranes were then imaged on a LI-COR Odyssey scanner with the 700 and 800 nm channels.

### Measurement of Piebaldism

While at CSHL, photographs of anesthetized or euthanized mice with piebaldism were obtained using a digital camera with a ruler alongside each animal to facilitate measurement of the surface area of the spotting, using the software program ImageJ (National Institute of Health, Public Domain, BSD-2). For those mice photographed during anesthesia, they were continually sedated with isoflurane, and many of these mice underwent CT scanning. Mice were weighed at the time of photography. Data was plotted using GraphPad Prism 9.5.1 (San Diego,CA, USA) for Macintosh IOS.

### Statistical Analysis

Chi squared analyses were preformed analyzing maternal effect lethality using a statistical threshold of 0.05. For data comparisons of two groups, two-tailed unpaired student’s t-test or multiple unpaired t-tests were used to calculate statistical significances between means using GraphPad Prism version 9.5.1(528) (San Diego, CA, USA). All t-tests performed herein utilized a significance threshold of 0.05. For western blotting and total protein assays Welch’s two sample t-test and two-way ANOVAs were performed using R-Studio version 1.4.1564 (Boston, MS, USA) for Macintosh IOS. Asterisks denote statistical significance marked by *p<0.05, **p<0.01, ***p<0.0001. Simple linear regression models were used to analyze differences between length of gestation and weights of dams using GraphPad Prism version 9.5.1(528) (San Diego, CA, USA). For analyzing embryo weight, embryo area and embryonic age nonlinear exponential regression models were performed using GraphPad Prism version 9.5.1(528) (San Diego, CA, USA).

### Data Analysis

GraphPad Prism software version 9.5.1(528) (San Diego, CA, USA), Microsoft Excel version 16.72 (Redmond, WA, USA) and Fiji by ImageJ (National Institute of Health, Public Domain, BSD-2) were used.

## Acknowledgements

GJL would like to thank the staff of the animal facility and/or transgenic core facility at CSHL (Leyi Li, Jodi Coblentz, Rachel Rubino, and Lisa Bianco) and IBR (Michael Parascando) for their assistance. Alison Sebold assisted with dissection and characterization of mouse vertebrae. We thank Dr. Goo Taeg Oh at Ewha Woman’s University (Seoul, Korea) for providing the original *Naa10^−/y^* mice to us.

## Supporting Information

**S1 Fig: Attempted generation of mNaa10 S37P knockin mice. A)** Schematic representation of the genomic locus of Naa10 and the design of the targeting vector. The primers used for sequencing of the construct are indicated by arrows. **B)** Schematic representation of the genomic locus after targeting and recombination with Flp (excision of neomycin selection cassette) and/or FLP (inversion of mini gene). **C)** Shown are the sequences of the Naa10 mini gene before and after inversion with FLP.

**S2 Fig: Naa10 expression in Naa10 knockdown mice.** Tissue was dissected from male C57BL/6NTac mice harboring the Naa10 mini gene in silenced (mini) or activated/S37P (inv) orientation as well as WT/ mice as control. Protein and RNA were isolated in parallel. **A)** Western Blot analyses for Naa10 and Naa15 from WT/, mini/ and inv/ mice. GAPDH was used as loading control. **B)** qPCR analyses of WT/, mini/ and inv/ mice for Naa10. The expression was normalized to WT/ for each tissue.

**S3 Fig: Further demonstration of Naa10 knockdown in Naa10 minigene and inverted minigene mice. A)** Western blotting of liver lysates. B) Western blotting of heart and liver lysates.

**S4 Fig: Quantitation of piebaldism, showing extensive variability. A)** The piebaldism in mice of various ages was measured. **B)** The piebaldism measured in mice of various ages was normalized against their body weight, by dividing the square millimeters by the weight in grams, yielding the metric of mm^2^/g.

**S5 Fig: There is no obvious correlation with piebaldism and the genotypes with age or weight. A)** Piebaldism in various mouse genotypes compared to age in days. **B)** Piebaldism plotted against weight in grams. **C)** mm2/wt plotted against age in days.

**S6 Fig: Calvaria (skull) Bone Density measured using computerized tomography (*CT) scanning.* A**) Male mice of various ages. **B)** Male mice of all ages. Mean, with standard deviation. *P<0.05, ****P<0.0001. **C)** Female mice of various ages plotted together. **D)** Females of all ages plotted together. Mean, with standard deviation; ns (not significant).

**S7 Fig: Femur Bone Density measured using computerized tomography (*CT) scanning.* (A)** Male mice of various ages. **(B)** Male mice of all ages. Mean, with standard deviation. *P<0.05, **P<0.01 **(C)** Female mice of various ages plotted together. **(D)** Females of all ages plotted together. Mean, with standard deviation. *P<0.05; ns (not significant).

**S8 Fig: Naa10 expression is not detected by Western blot in mice bearing indels in *NAA10*.**

**S9 Fig. Quantification of *Naa10^−/Y^* and *Naa10^+/Y^* heart and liver tissue lysate. A)** Membranes incubated in rabbit anti-NAA10 MAb and goat-anti-rabbit secondary (800 nm channel). Biological replicates (n = 8) were obtained of *Naa10^−/Y^*(N = 4) and *Naa10^+/Y^* (N = 4) mice. Heart and tissue lysate were obtained from each mouse. Blots were stained for total protein (REVERT 700 Total protein stain) post-transfer. After stain removal, blots were incubated in anti-NAA10 MAb and anti-rabbit secondary antibody. NAA10 signal was normalized to total protein. **B)** Membranes stained for total protein using REVERT 700 total protein stain (700 nm channel) to verify transfer and equal loading for NAA10 signal normalization. **C)** Quantification of normalized NAA10 signal in heart and liver lysate; horizontal crossbar indicates mean (± SD; 2-way ANOVA, F statistic = 14.52 on 3 and 12 DF, *P < 0.05)

**S10 Fig. NAA10 immunoblotting in heterozygous females.** Whole membranes corresponding to representative immunoblots of liver and heart lysates in Figure 1. Membranes were stained for total protein after transfer using REVERT 700 total protein stain. After total protein stain removal, membranes were incubated in rabbit anti-NAA10 MAb and goat-anti-rabbit secondary antibodies. From top to bottom, target protein (NAA10) and loading control (total protein). **A)** Representative blot for immunoblotted liver lysate. From top to bottom, NAA10 excerpt, NAA10 whole membrane, and total protein stained membrane. **B)** Representative blot for immunoblotted heart lysate. From top to bottom, NAA10 excerpt, NAA10 whole membrane, and total protein stained membrane.

**S11 Fig. Western blot analysis of NAA10 signal in heterozygous females and C57 controls.** Liver lysates were obtained from *Naa10^−/+^*mice (N = 3) and C57BL/6J females (N = 3). *Naa10^+/Y^* lysates were loaded to balance gels and excluded from analysis. Both replicates are shown. Blots were stained for total protein post-transfer and scanned to verify successful transfer of equal loading for use as loading control; after total protein stain removal, blots were incubated with anti-NAA10 MAb and anti-rabbit secondary. **A-B)** NAA10 lanes and whole membranes from replicate blots incubated with anti-NAA10 Mab and goat-anti-rabbit secondary antibody. Immunostained membranes were scanned in 800nm channel. **C)** Whole membrane after post-transfer staining for total membrane using REVERT 700 total protein stain to verify transfer and equal loading for NAA10 signal normalization. Total-protein-stained membranes were scanned in 700nm channel. **D)** Quantification of NAA10 signal normalized to total protein. Short black crossbar indicates mean NAA10 signal normalized to total protein (±SD, Welch’s 2-sample t-test, t = −1.6784, df = 9.1762, P = 0.1252).

**S12 Fig.** Neonatal phenotypes of *Naa10* knockout mice after 20 backcrosses**. A)** Pictures of *Naa10^−/Y^* (left) *and Naa10^+/Y^* (right) litter mates. The *Naa10^−/Y^* mouse is noticeably smaller in size and body/skull composition. **B)** Piebaldism present on ventral abdomen of *Naa10^−/Y^* mouse (left) compared to *Naa10^+/Y^* (right) litter mate. **C)** Hydronephrosis of the left kidney in a separate *Naa10^−/Y^*.

**S1 Table. Genotypes from female *Naa10^mini/WT^* female mice crossed to *C57BL/6J* male mice, after weaning.**

**S2 Table. Genotypes from female *Naa10^inv/WT^* female mice crossed to *C57BL/6J* male mice, after weaning.**

**S3 Table. Skeletal abnormalities of ribs and sternebrae in mice with varying genotypes. S4 Table. Cervical Vertebrae Fusions**

**S5 Table. Zygotes injected during attempts to generate Naa10 Ser37Pro mutant mice**

**S6 Table. Genotypes from female *Naa10*^Δ668*/WT*^ female mice crossed to *Naa10*^Δ668*/y*^ male mice, post weaning.**

**S7 Table. Genotypes from female *Naa10*^Δ668-674*/WT*^ female mice crossed to *C57BL/6J* male mice, after weaning.**

**S8 Table**: **Primers used for genotyping or RT-PCR**:

**S9 Table: Detailed information for antibodies used in western blotting.**

