## Supplementary Information for "Evaluating possible maternal effect lethality and genetic background effects in *Naa10* knockout mice"

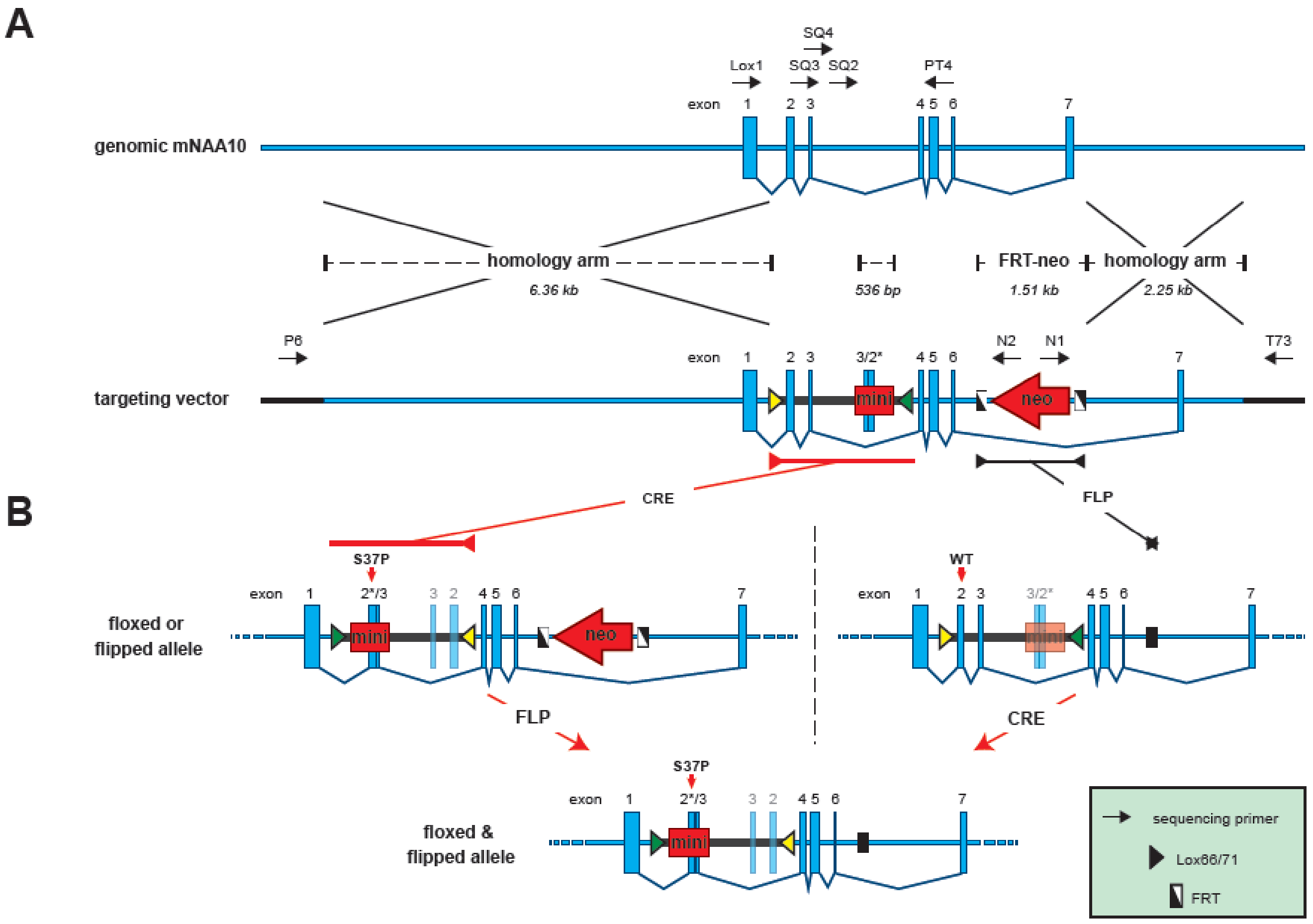


C

Naa10 mini-gene 5'>3': intron1-Exon2 with **point mutation**-Exon3-intron3
ACTCAGTCCTGGAACTGGCACGTTTTGCATGCCGCATGGAAACAGGGCTTGTGCAAAGTCGTATAGTAGCAACTGTGCCCTGCCCC
GGGGAAAGGGGAGACCCAGAGATCGACGTGTGTGGCCCTCTTAGTCATTGCAGCCCTGGACTAACCGTATCCCACTCTCATGCAG
CCCGAAGACCTGATGAACATGCAGCACTGCAACCTTCTCTGCCTGCCGGAGAACTACCAGATGAAGTACTATTTCTATCATGGCCTC
**C**CTTGGCCCCAGCTTTCTTACATTGCTGAGGATGAGAATGGGAAGATTGTGGGCTACGTCTTGGCTAAAATGTGAGTCACCAAGGA
GAAAGAAAAGACACTAATGTTAAGAAGGACAAAAGCTTTGGAGAAGGGAATGGGAAGCGGTGAATTAGTTTGAGAGGCATTCTCTGT
TGTCCAGATTGGTGCTCACTCAGGCTAACCTCAAACTTATAGTAGTCTTCATGCTGAGCC

Naa10 mini-gene 3'>5': intron3-Exon3-Exon2 **with point mutation**-intron1
GGCTCAGCATGAAGACTACTATAAGTTTGAGGTTAGCCTGAGTGAGCACCAATCTGGACAACAGAGAATGCCTCTCAAACTAATTCA
CCGCTTCCCATTCCCTTCTCCAAAGCTTTTGTCCTTCTTAACATTAGTGTCTTTTCTTTCTCCTTGGTGACTCACATTTTAGCCAAGAC
GTAGCCCACAATCTTCCCATTCTCATCCTCAGCAATGTAAGAAAGCTGGGGCCAAG**G**GAGGCCATGATAGAAATAGTACTTCATCTG
GTAGTTCTCCGGCAGGCAGAGAAGGTTGCAGTGCTGCATGTTCATCAGGTCTTCGGGCTGCATGAGAGTGGGATACGGTTAGTCCA
GGGCTGCAATGACTAAGAGGGCCACACACGTCGATCTCTGGGTCTCCCCTTTCCCCGGGGCAGGGCACAGTTGCTACTATACGAC
TTTGCACAAGCCCTGTTTCCATGCGGCATGCAAAACGTGCCAGTTCCAGGACTGAGT

**S1 Fig: Attempted generation of mNaa10 S37P knockin mice. A)** Schematic representation of the genomic locus of Naa10 and the design of the targeting vector. The primers used for sequencing of the construct are indicated by arrows. **B)** Schematic representation of the genomic locus after targeting and recombination with Flp (excision of neomycin selection cassette) and/or FLP (inversion of mini gene). **C)** Shown are the sequences of the Naa10 mini gene before and after inversion with FLP.

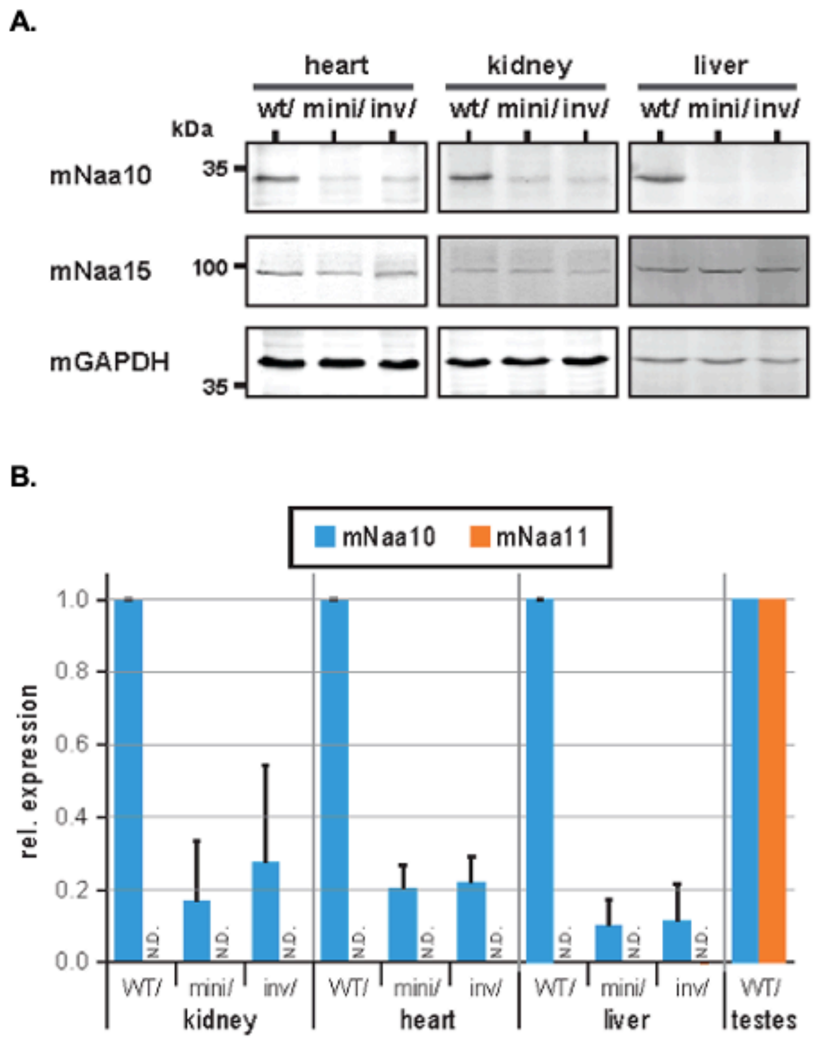


**S2 Fig: Naa10 expression in Naa10 knockdown mice.** Tissue was dissected from male C57BL/6NTac mice harboring the Naa10 mini gene in silenced (mini) or activated/S37P (inv) orientation as well as WT/ mice as control. Protein and RNA were isolated in parallel. **A)** Western Blot analyses for Naa10 and Naa15 from WT/, mini/ and inv/ mice. GAPDH was used as loading control. **B)** qPCR analyses of WT/, mini/ and inv/ mice for Naa10. The expression was normalized to WT/ for each tissue.

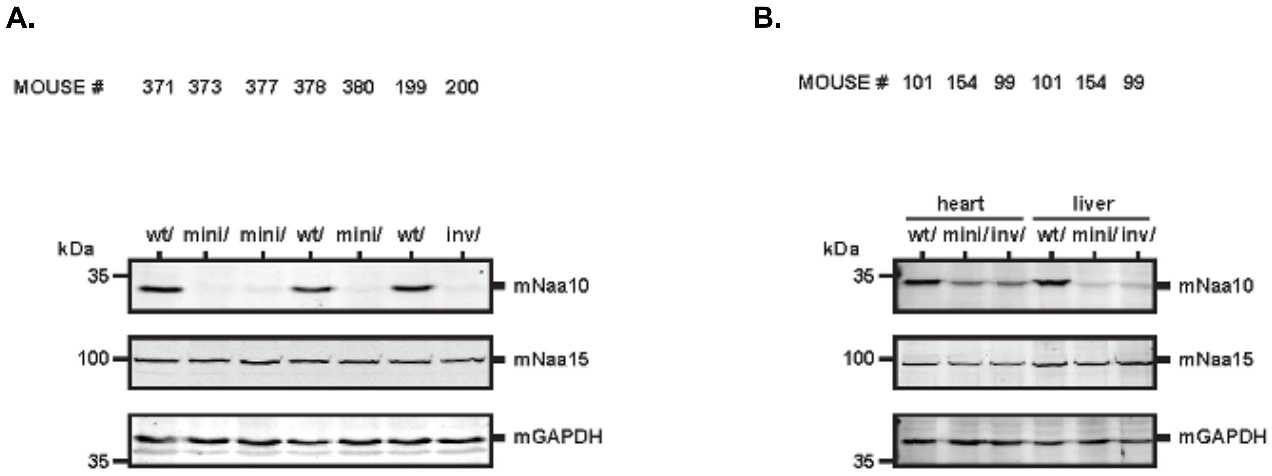


**S3 Fig: Further demonstration of Naa10 knockdown in Naa10 minigene and inverted minigene mice. A)** Western blotting of liver lysates. B) Western blotting of heart and liver lysates.

A.

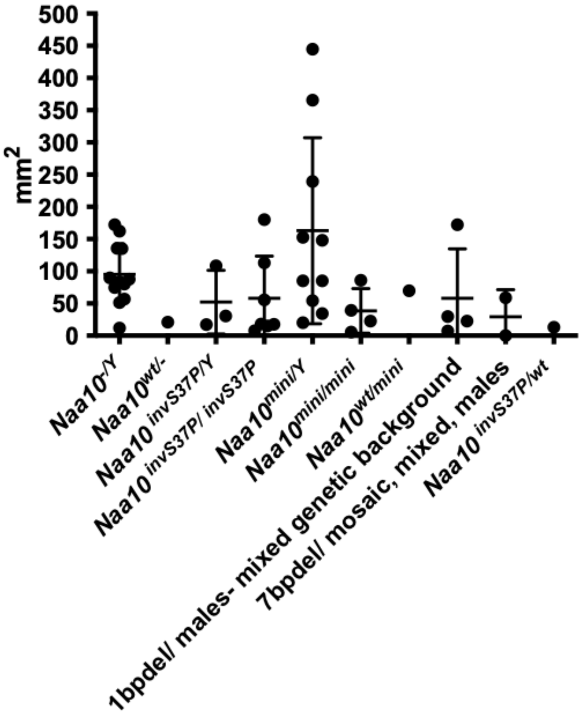


B.

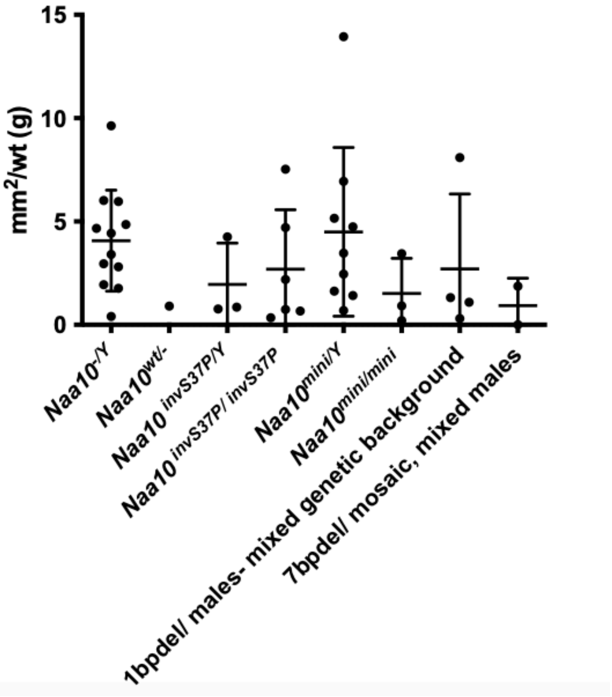


**S4 Fig: Quantitation of piebaldism, showing extensive variability. A)** The piebaldism in mice of various ages was measured. **B)** The piebaldism measured in mice of various ages was normalized against their body weight, by dividing the square millimeters by the weight in grams, yielding the metric of $mm^2/g$.

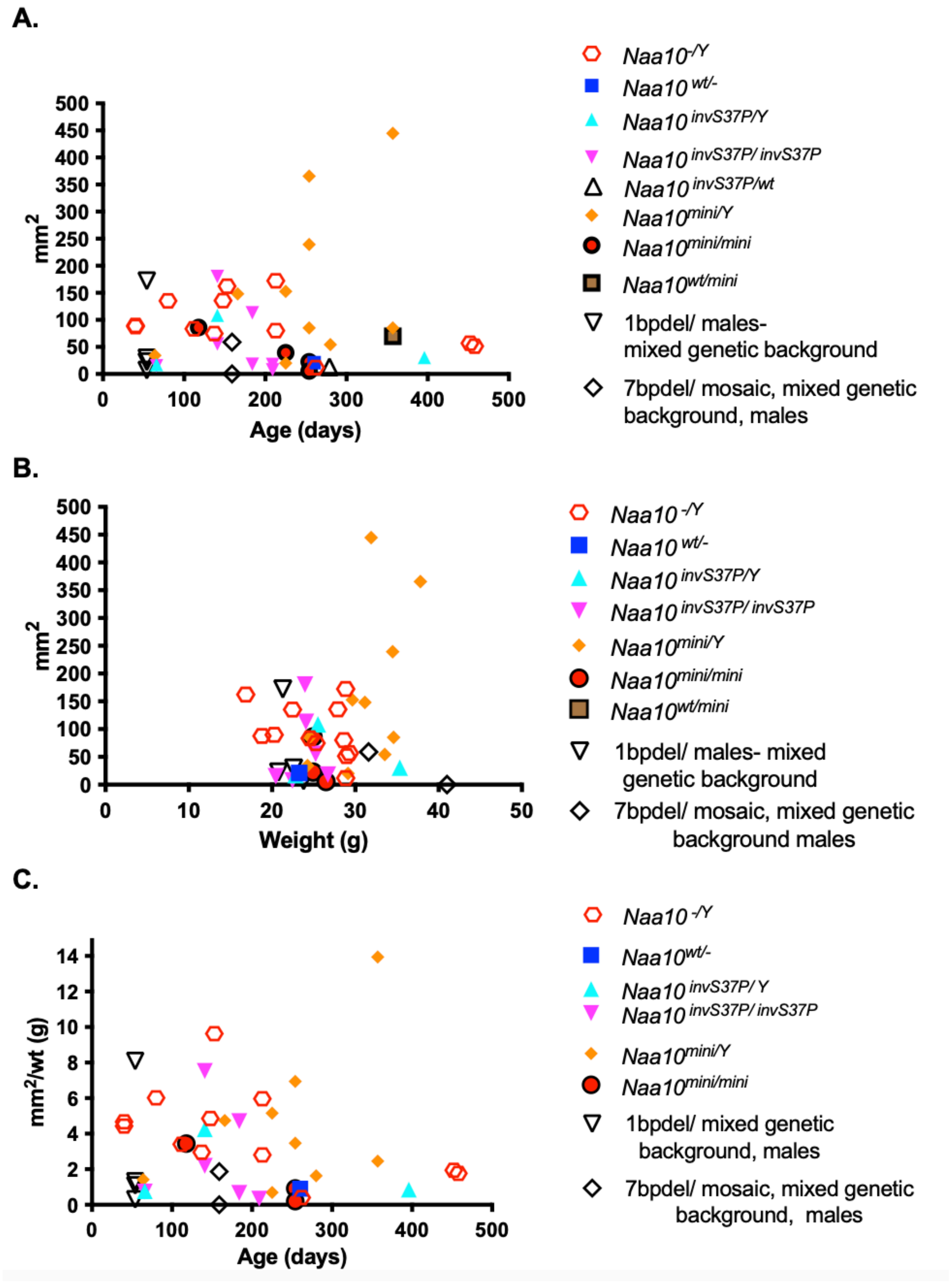


**S5 Fig: There is no obvious correlation with piebaldism and the genotypes with age or weight. A)** Piebaldism in various mouse genotypes compared to age in days. **B)** Piebaldism plotted against weight in grams. **C)** mm2/wt plotted against age in days.

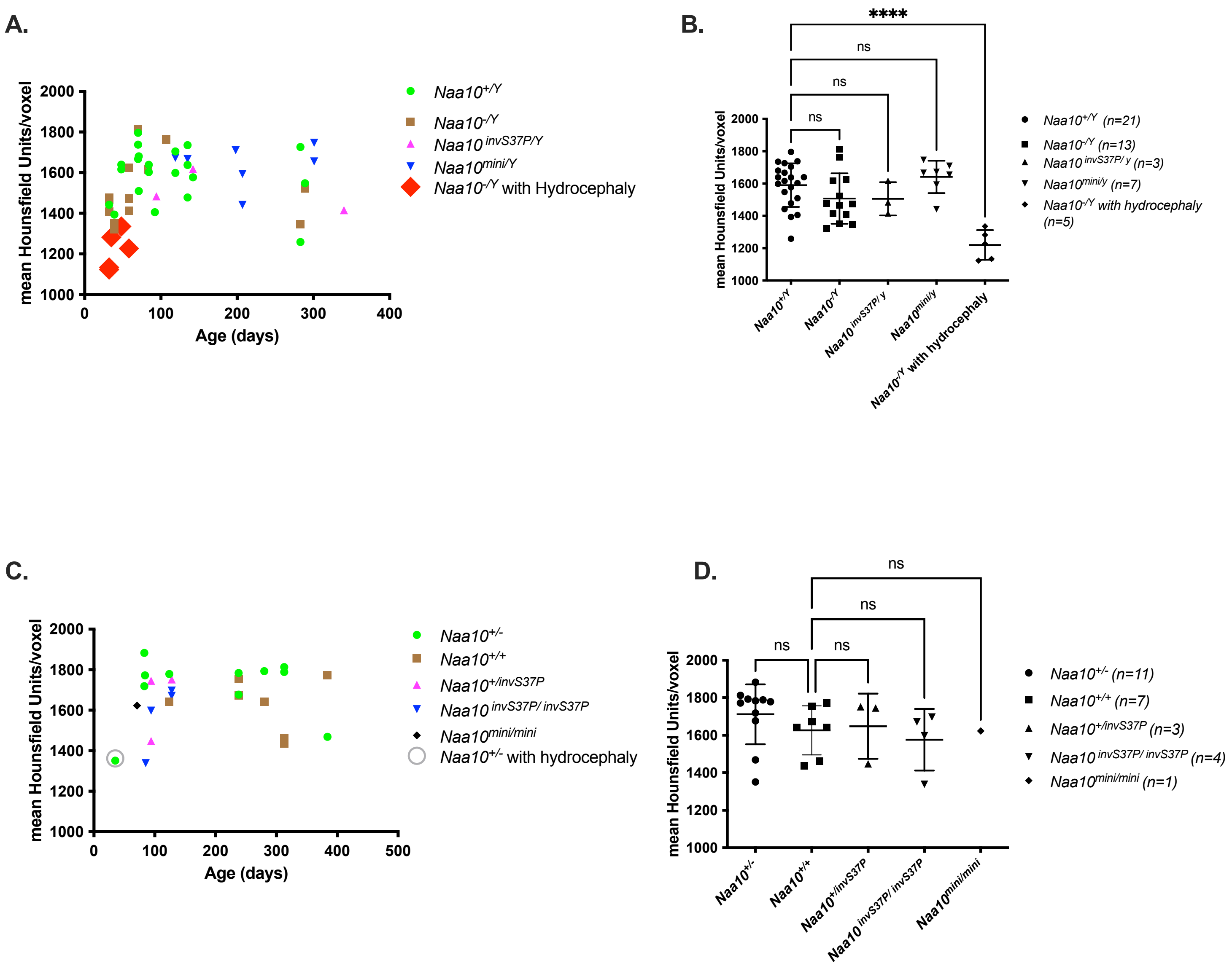


**S6 Fig: Calvaria (skull) Bone Density measured using computerized tomography (*CT)* scanning. A**) Male mice of various ages. **B)** Male mice of all ages. Mean, with standard deviation. *P<0.05, ****P<0.0001. **C)** Female mice of various ages plotted together. **D)** Females of all ages plotted together. Mean, with standard deviation; ns (not significant).

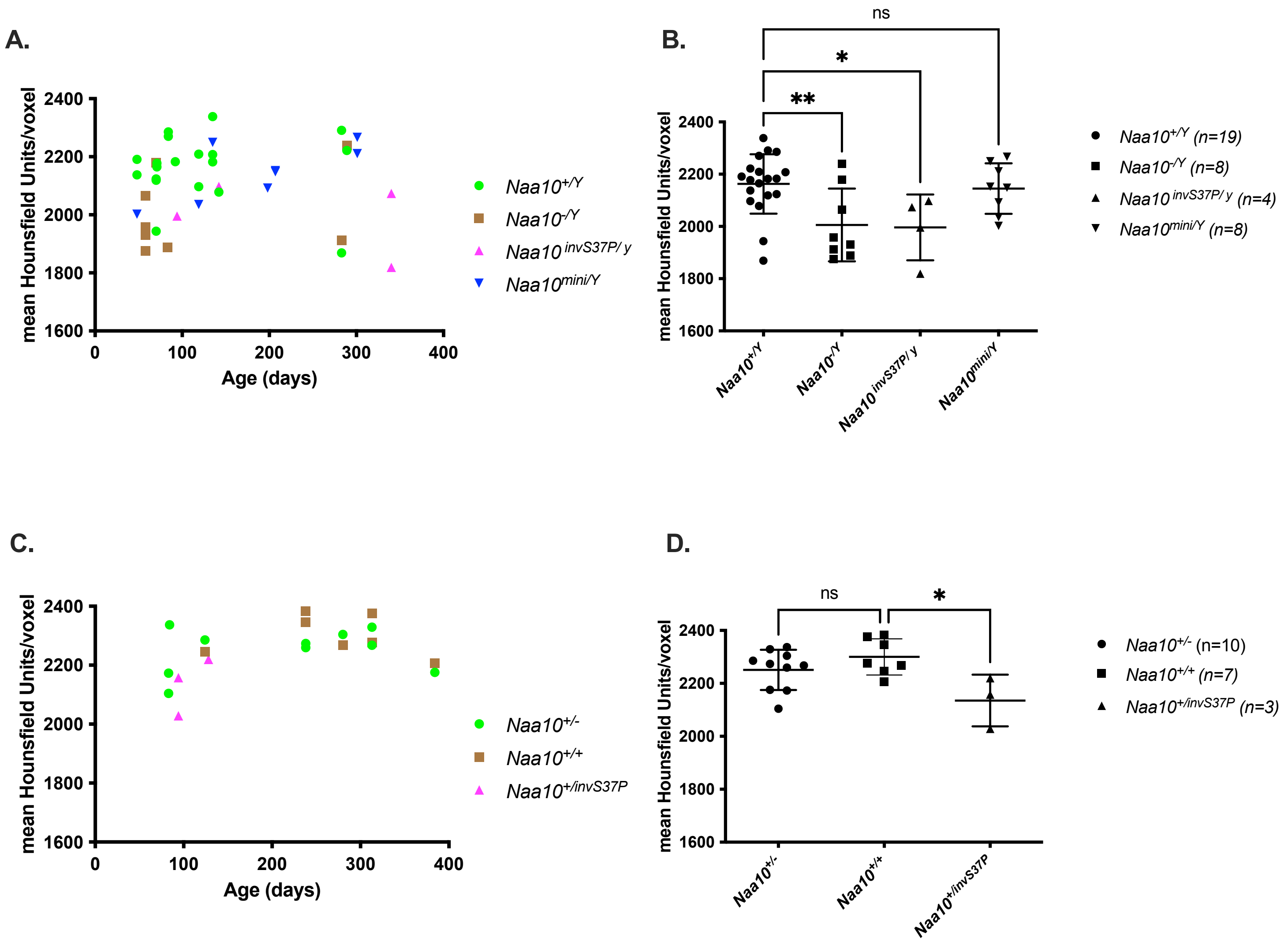


**S7 Fig: Femur Bone Density measured using computerized tomography (*CT)* scanning. (A)** Male mice of various ages. **(B)** Male mice of all ages. Mean, with standard deviation. *P<0.05, **P<0.01 **(C)** Female mice of various ages plotted together. **(D)** Females of all ages plotted together. Mean, with standard deviation. *P<0.05; ns (not significant).

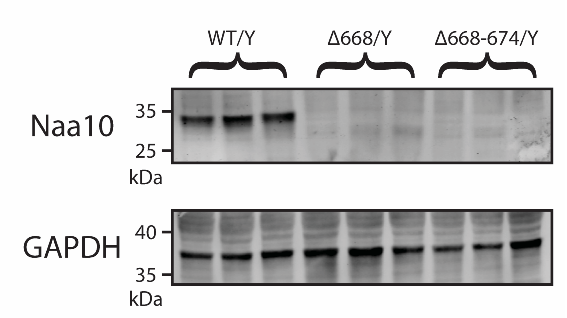


**S8 Fig: Naa10 expression is not detected by Western blot in mice bearing indels in *NAA10*.**

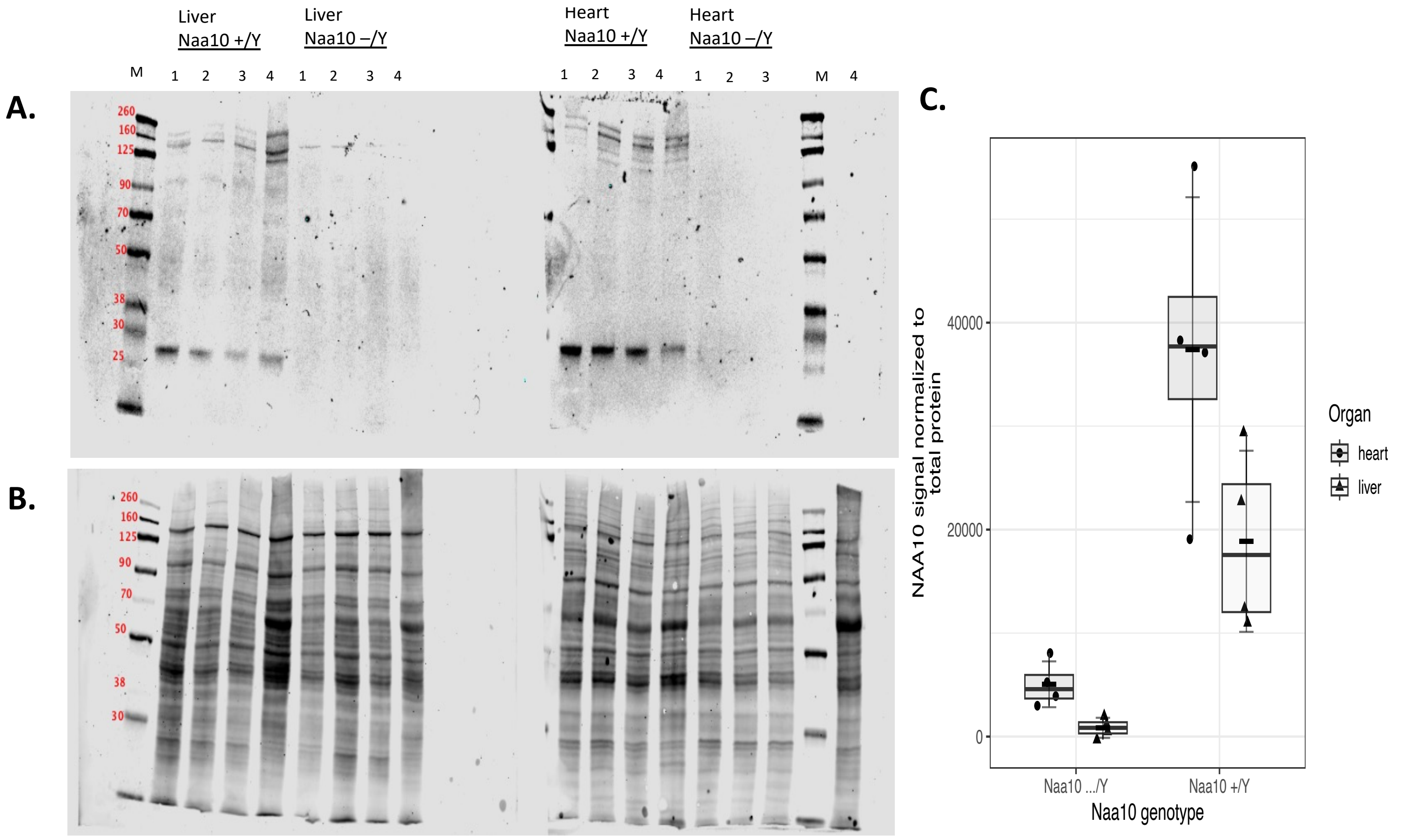


**S9 Fig. Quantification of *Naa10*$^{-/Y}$ and *Naa10*$^{+/Y}$ heart and liver tissue lysate.**
**A)** Membranes incubated in rabbit anti-NAA10 MAb and goat-anti-rabbit secondary (800 nm channel). Biological replicates (n = 8) were obtained of *Naa10*$^{-/Y}$ (N = 4) and *Naa10*$^{+/Y}$ (N = 4) mice. Heart and tissue lysate were obtained from each mouse. Blots were stained for total protein (REVERT 700 Total protein stain) post-transfer. After stain removal, blots were incubated in anti-NAA10 MAb and anti-rabbit secondary antibody. NAA10 signal was normalized to total protein. **B)** Membranes stained for total protein using REVERT 700 total protein stain (700 nm channel) to verify transfer and equal loading for NAA10 signal normalization. **C)** Quantification of normalized NAA10 signal in heart and liver lysate; horizontal crossbar indicates mean (± SD; 2-way ANOVA, F statistic = 14.52 on 3 and 12 DF, *$P < 0.05$)

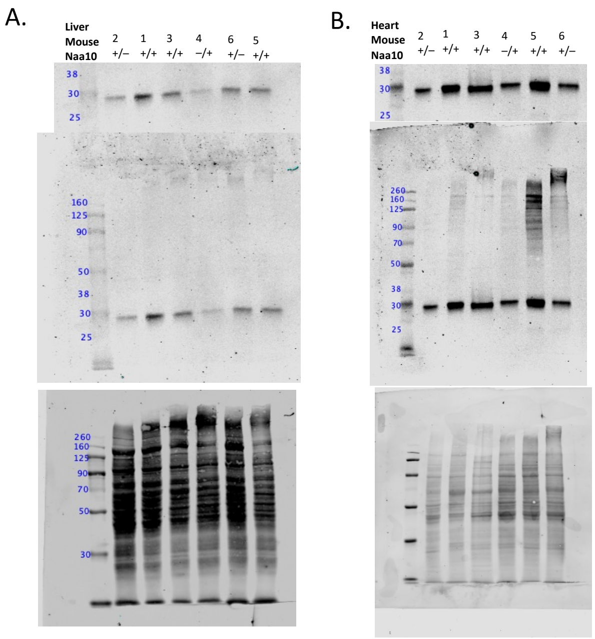


**S10 Fig. NAA10 immunoblotting in heterozygous females.** Whole membranes corresponding to representative immunoblots of liver and heart lysates in Figure 1. Membranes were stained for total protein after transfer using REVERT 700 total protein stain. After total protein stain removal, membranes were incubated in rabbit anti-NAA10 MAb and goat-anti-rabbit secondary antibodies. From top to bottom, target protein (NAA10) and loading control (total protein). **A)** Representative blot for immunoblotted liver lysate. From top to bottom, NAA10 excerpt, NAA10 whole membrane, and total protein stained membrane. **B)** Representative blot for immunoblotted heart lysate. From top to bottom, NAA10 excerpt, NAA10 whole membrane, and total protein stained membrane.

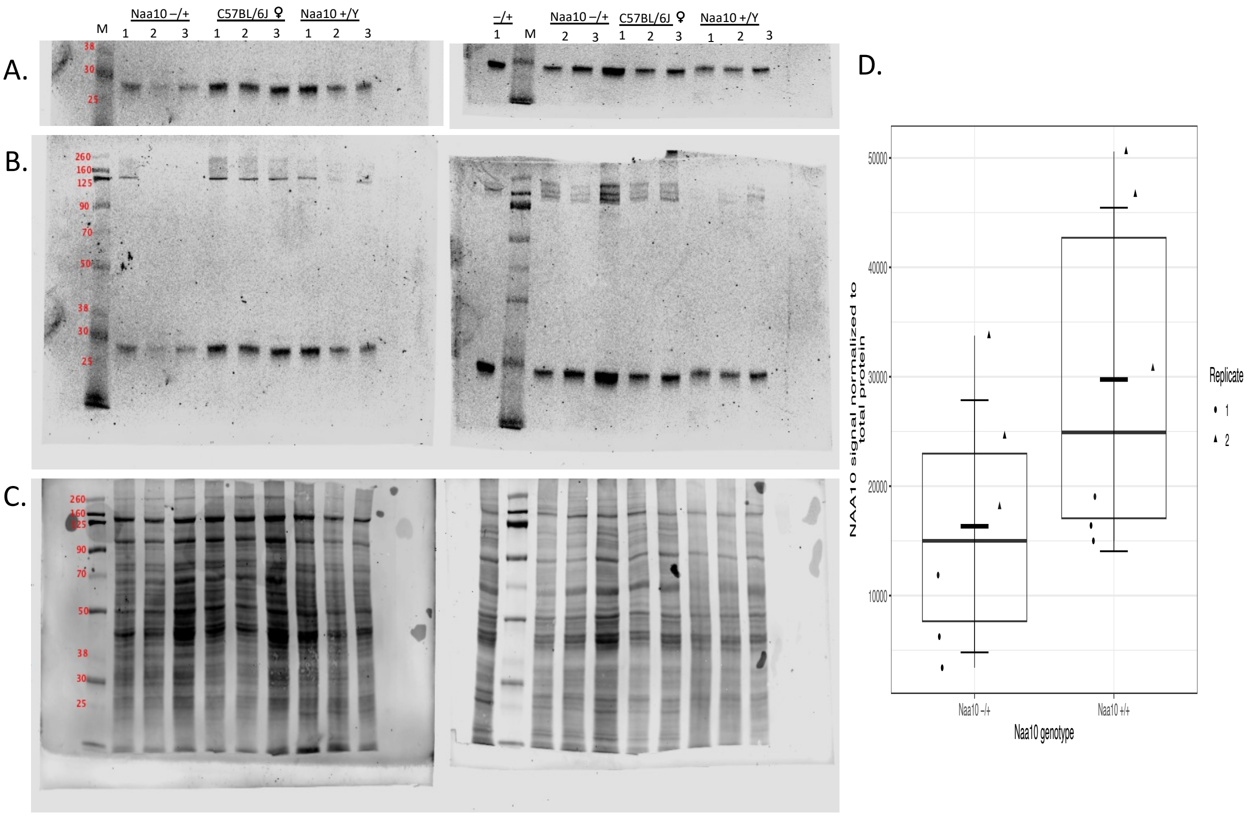


**S11 Fig. Western blot analysis of NAA10 signal in heterozygous females and C57 controls.** Liver lysates were obtained from *Naa10*$^{-/+}$ mice (N = 3) and C57BL/6J females (N = 3). *Naa10*$^{+/Y}$ lysates were loaded to balance gels and excluded from analysis. Both replicates are shown. Blots were stained for total protein post-transfer and scanned to verify successful transfer of equal loading for use as loading control; after total protein stain removal, blots were incubated with anti-NAA10 MAb and anti-rabbit secondary. **A-B)** NAA10 lanes and whole membranes from replicate blots incubated with anti-NAA10 Mab and goat-anti-rabbit secondary antibody. Immunostained membranes were scanned in 800nm channel. **C)** Whole membrane after post-transfer staining for total membrane using REVERT 700 total protein stain to verify transfer and equal loading for NAA10 signal normalization. Total-protein-stained membranes were scanned in 700nm channel. **D)** Quantification of NAA10 signal normalized to total protein. Short black crossbar indicates mean NAA10 signal normalized to total protein (±SD, Welch's 2-sample t-test, t = -1.6784, df = 9.1762, P = 0.1252).

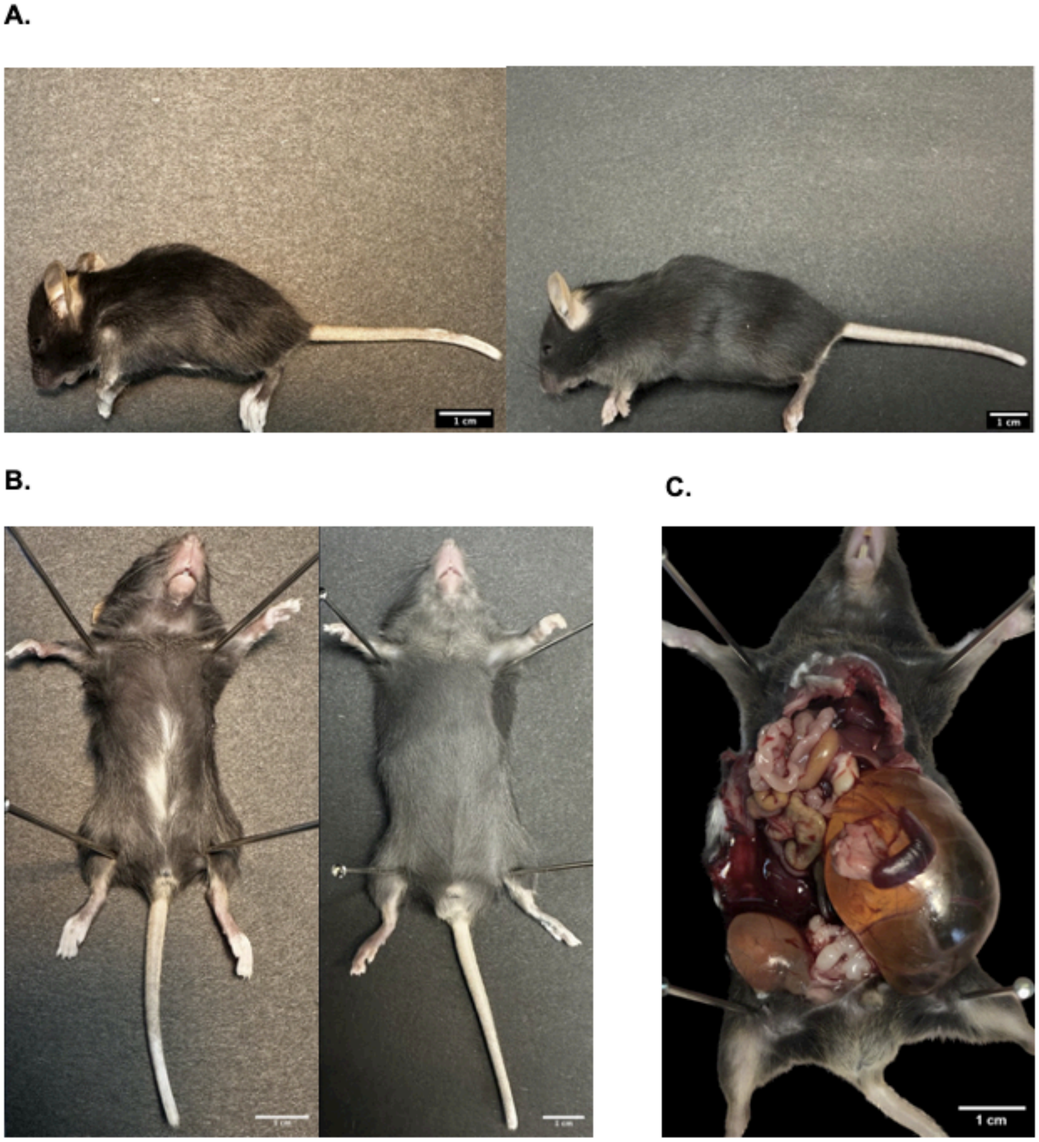


**S12 Fig.** Neonatal phenotypes of *Naa10* knockout mice after 20 backcrosses**. A)** Pictures of *Naa10*$^{-/Y}$ (left) *and Naa10*$^{+/Y}$ (right) litter mates. The *Naa10*$^{-/Y}$ mouse is noticeably smaller in size and body/skull composition. **B)** Piebaldism present on ventral abdomen of *Naa10*$^{-/Y}$ mouse (left) compared to *Naa10*$^{+/Y}$ (right) litter mate. **C)** Hydronephrosis of the left kidney in a separate *Naa10*$^{-/Y}$.

**S1 Table. Genotypes from female *Naa10*$^{mini/WT}$ female mice crossed to *C57BL/6J* male mice, after weaning.**

| Genotype (Expected Mendelian %) | Naa10$^{+/y}$ (25%) | Naa10$^{mini/y}$ (25%) | Naa10$^{+/+}$ (25%) | Naa10$^{mini/+}$ (25%) |
|---|---|---|---|---|
| **Adults (n=74)** | 18 (24%) | 16 (22%) | 22 (30%) | 18 (24%) |

Expected and observed Mendelian genotype ratios in offspring from crosses.

**S2 Table. Genotypes from female *Naa10*$^{inv/WT}$ female mice crossed to *C57BL/6J* male mice, after weaning.**

| Genotype (Expected Mendelian %) | Naa10$^{+/y}$ (25%) | Naa10$^{inv/y}$ (25%) | Naa10$^{+/+}$ (25%) | Naa10$^{inv/+}$ (25%) | Unable to genotype |
|---|---|---|---|---|---|
| **Adults (n=106)** | 27 (25%) | 13 (12%) | 37 (35%) | 26 (25%) | 3 (3%) |

Expected and observed Mendelian genotypes ratios in offspring from crosses.

### S3 Table. Skeletal abnormalities of ribs and sternebrae in mice with varying genotypes.

| | **WT/inv (n=4)** | **WT/mini (n=1)** | **inv/inv (n=7)** | **mini/mini (n=2)** | **WT/1bp (n=3)** | **1bp/1bp (n=5)** | **mini/Y (n=17)** | **inv/Y (n=10)** | **1bp/Y (n=20)** | **7bp/Y (n=4)** | **7bp/Y MOSAIC (n=2)** |
|---|---|---|---|---|---|---|---|---|---|---|---|
| 4 sternebrae | 2(50,0%) | 0(0%) | 4(57.1%) | 1(50%) | 1(33.3%) | 3(60%) | 13(76.5%) | 8(80%) | 13(65%) | 3(75%) | 1 (50%) |
| 3 sternebrae | 2(50,0%) | 1(100%) | 0(0%) | 1(50%) | 1(33.3%) | 1(20%) | 3(17,6%) | 0(0%) | 1(05%) | 0(0%) | 0(0%) |
| 4 sternebrae but with 3/4 fusion | 0(0%) | 0(0%) | 3(42,9%) | 0(0%) | 1(33.3%) | 1(20%) | 1(5,9%) | 2(20,0%) | 6(30%) | 1(25%) | 1 (50%) |
| 14 ribs total bilaterally | 0(0%) | 0(0%) | 7(100,0%) | 2(100,0%) | 0(0,0%) | 5(100%) | 17(100%) | 10(100%) | 20(100%) | 4(100%) | 1 (50%) |
| 13 ribs total bilaterally | 4(100%) | 1(100%) | 0(0%) | 0(0%) | 3(100%) | 0(0%) | 0(0%) | 0(0%) | 0(0%) | 0(0%) | 1 (50%) |
| 8 ribs attached to sternum bilaterally | 0(0%) | 0(0%) | 7(100,0%) | 2(100,0%) | 0(0%) | 4(80%) | 17(100%) | 10(100%) | 18(90%) | 4(100%) | 0(0%) |
| 7 ribs attached to sternum bilaterally | 3(75%) | 1(100%) | 0(0%) | 0(0%) | 3(100%) | 0(0%) | 0(0%) | 0(0%) | 2(10%) | 0(0%) | 1 (50%) |
| 7 ribs linking to sternum on one side, 8 on the other side | 1(25%) | 0(0%) | 0(0%) | 0(0%) | 0(0%) | 1(20%) | 0(0%) | 0(0%) | 0(0%) | 0(0%) | 1 (50%) |
| 14 Thoracic vertebrae | 0(0%) | 0(0%) | 7(100%) | 2(100%) | 0(0,0%) | 5(100%) | 17(100,0%) | 10(100%) | 18(90%) | 4(100%) | 1 (50%) |
| 13 Thoracic Vertebrae | 4(100%) | 1(100%) | 0(0%) | 0(0%) | 3(100%) | 0(0%) | 0(0%) | 0(0%) | 2(10%) | 0(0%) | 1 (50%) |
| Total | 16 | 4 | 28 | 8 | 12 | 20 | 68 | 40 | 68 | 16 | 8 |

### S4 Table. Cervical Vertebrae Fusions

| | wt/inv | wt/mini | inv/ | inv/inv | mini/ | mini/mini | ko/ | ko/ko | 7bp/ | 1bp/ | 1bp/1bp |
|---|---|---|---|---|---|---|---|---|---|---|---|
| **One or more fusion events** | 3/3 (100%) | 1/1 (100%) | 8/9 (89%) | 6/6 (100%) | 11/11 (100%) | 4/4 (100%) | 9/10 (90%) | 1/1 (100%) | 5/5 (100%) | 1/2 (50%) | 1/1 (100%) |
| **Two or more fusion events** | 3/3 (100%) | 1/1 (100%) | 4/9 (44%) | 5/6 (83%) | 5/11 (45%) | 3/4 (75%) | 3/9 (33%) | 1/1 (100%) | 2/4 (50%) | 0/2 (0%) | 0/1 (0%) |
| **Consecutive fusion events** | 2/3 (67%) | 0/1 (0%) | 3/8 (38%) | 4/6 (67%) | 4/11 (36%) | 2/4 (50%) | 1/9 (11%) | 1/1 (100%) | 2/4 (50%) | 0/2 (0%) | 0/1 (0%) |
| **C1+2 fusion** | 2/3 (67%) | 1/1 (100%) | 8/9 (89%) | 5/5 (100%) | 11/11 (100%) | 4/4 (100%) | 7/10 (70%) | 1/1 (100%) | 4/4 (100%) | 1/2 (50%) | 0/1 (0%) |
| **C2+3 fusion** | 2/3 (67%) | 0/1 (0%) | 1/8 (13%) | 2/5 (40%) | 4/11 (36%) | 1/4 (25%) | 2/9 (22%) | 1/1 (100%) | 2/4 (50%) | 0/2 (0%) | 1/1 (100%) |
| **C3+4 fusion** | 1/3 (33%) | 1/1 (100%) | 0/7 (0%) | 1/5 (20%) | 2/11 (18%) | 1/4 (25%) | 2/9 (22%) | 0/1 (0%) | 2/5 (40%) | 0/2 (0%) | 0/1 (0%) |
| **C4+5 fusion** | 1/3 (33%) | 0/1 (0%) | 1/9 (11%) | 0/5 (0%) | 1/11 (9%) | 2/4 (50%) | 1/9 (11%) | 0/1 (0%) | 0/6 (0%) | 0/2 (0%) | 0/1 (0%) |
| **C5+6 fusion** | 0/3 (0%) | 0/1 (0%) | 1/8 (13%) | 0/5 (0%) | 1/11 (9%) | 1/4 (25%) | 0/9 (0%) | 0/1 (0%) | 0/6 (0%) | 0/2 (0%) | 0/1 (0%) |
| **C6+7 fusion** | 0/3 (0%) | 0/1 (0%) | 3/9 (33%) | 1/6 (17%) | 1/11 (9%) | 0/4 (0%) | 0/9 (0%) | 0/1 (0%) | 1/6 (17%) | 0/2 (0%) | 0/1 (0%) |
| **C7+ T1 fusion** | 1/3 (33%) | 0/1 (0%) | 1/9 (11%) | 4/6 (67%) | 2/11 (18%) | 2/4 (50%) | 1/9 (11%) | 0/1 (0%) | 0/6 (0%) | 0/2 (0%) | 0/1 (0%) |
| **T1+2 fusion** | 0/3 (0%) | 0/1 (0%) | 2/9 (22%) | 2/6 (33%) | 0/11 (0%) | 1/4 (25%) | 0/9 (0%) | 0/1 (0%) | 0/6 (0%) | 0/2 (0%) | 0/1 (0%) |
| **Total** | 3 | 1 | 9* | 6* | 12* | 4 | 10* | 1 | 6* | 5* | 1 |

Due to prior loss or damage of vertebrae in some samples, n for each specific fusion event was adjusted according to the number of vertebrae available for examination.

### S5 Table. Zygotes injected during attempts to generate Naa10 Ser37Pro mutant mice

| Date | Construct | Cas9 mRNA ng/ul | Cas9 protein (ng/ul) | Donor DNA (ng/ul) | Zygote injected | Intact | Pups DOB | Pups number |
|---|---|---|---|---|---|---|---|---|
| 6/7/2016 | NAA10 | 50 | | 100 | 147 | 98 | 6/25/16-6/26/16 | 43 |
| 6/9/2016 | NAA10 | 50 | | 100 | 81 | 73 | 6/28/2016 | 32 |
| 9/9/2016 | NAA10 | | 50 | 50 | 76 | 72 | 9/28/2016 | 24 |
| 12/13/2016 | NAA10 | | 50 | 100 | 134 | 120 | 12/30-12/31/2016 | 50 |
| 12/15/2016 | NAA10 | | 50 | 50 | 100 | 60 | 1/2/2017 | 7 |

### S6 Table. Genotypes from female *Naa10*$^{\Delta 668/WT}$ female mice crossed to *Naa10*$^{\Delta 668/y}$ male mice, post weaning.

| Genotype (Expected Mendelian %) | Naa10$^{+/y}$ (25%) | Naa10$^{\Delta 668/y}$ (25%) | Naa10$^{\Delta 668/WT}$ (25%) | Naa10$^{\Delta 668/\Delta 668}$ (25%) | Unable to genotype |
|---|---|---|---|---|---|
| **Adults (n=174)** | 41 (24%) | 59 (34%) | 41 (24%) | 32 (18%) | 1 (1%) |

Expected and observed Mendelian ratio of genotypes in offspring from crosses.

**S7 Table. Genotypes from female *Naa10*$^{\Delta 668\text{-}674/WT}$ female mice crossed to *C57BL/6J* male mice, after weaning.**

| Genotype (Expected Mendelian %) | *Naa10*$^{+/y}$ (25%) | *Naa10*$^{\Delta 668\text{-}674/y}$ (25%) | *Naa10*$^{+/+}$ (25%) | *Naa10*$^{\Delta 668\text{-}674/WT}$ (25%) | Unable to genotype |
|---|---|---|---|---|---|
| **Adults (n=87)** | 26 (30%) | 22 (25%) | 21 (24%) | 15 (17%) | 3 (3%) |

Expected and observed Mendelian ratio of genotypes in offspring from crosses.

**S8 Table**: **Primers used for genotyping or RT-PCR**:

| Gene | Forward primer | Reverse primer |
|---|---|---|
| mNaa10-Exon2/3 | ctcttggccccagctttctt | |
| mNaa10-Exon3/4 | | tcgtctgggtcctcttccat |
| mNaa11 | accccacaagcaaagacagtg | agcgatgctcaggaaatgctct |
| mGAPDH | aggtcggtgtgaacggatttg | tgtagaccatgtagttgaggtca |
| mACTB_F | ggctgtattcccctccatcg | ccagttggtaacaatgccatgt |

**S9 Table: Detailed information for antibodies used in western blotting.**

| Target | Manufacturer (Catalog no, City, State) | Epitope/Immunogen |
|---|---|---|
| Anti-Naa10/ARDA1A (Rabbit polyclonal) | Abcam (Cat# ab155687, Boston, MA) | Recombinant fragment, corresponding to a region within amino acids 1-235 of Human ARD1A (UniProt: P41227). |
| Anti-Naa10/ARDA1A (Rabbit monoclonal) | Cell Signaling (Cat# 13357, Danvers, MA) | Monoclonal antibody is produced by immunizing animals with a synthetic peptide corresponding to residues surrounding Asp204 of human ARD1A protein. |
| Anti-Naa15/NARG1 (Mouse monoclonal-IGg1) | Abcam (Cat# ab60065, Boston, MA) | Recombinant fragment (Human) |
| Anti-GAPDH (Mouse monoclonal-IGg2b) | Abcam (Cat# ab9484, Boston, MA) | |
